# Melbournevirus-encoded histone doublets are recruited to virus particles and form destabilized nucleosome-like structures

**DOI:** 10.1101/2021.04.29.441998

**Authors:** Yang Liu, Chelsea Marie Toner, Nadege Philippe, Sandra Jeudy, Keda Zhou, Samuel Bowerman, Alison White, Garrett Edwards, Chantal Abergel, Karolin Luger

**Affiliations:** Department of Biochemistry, University of Colorado at Boulder, 80309 Boulder, Colorado; Howard Hughes Medical Institute; Aix–Marseille University, Centre National de la Recherche Scientifique, Information Génomique & Structurale, Unité Mixte de Recherche 7256 (Institut de Microbiologie de la Méditerranée, FR3479, IM2B), 13288 Marseille Cedex 9, France

**Keywords:** NCLDV, giant virus, viral nucleosome, doublet histone, cryoEM, viral factory, histone tail, acidic patch, non-eukaryotic nucleosome, nucleosome-like-particle

## Abstract

The organization of genomic DNA into defined nucleosomes has long been viewed as a hallmark of eukaryotes. This paradigm has been challenged by the identification of ‘minimalist’ histones in archaea, and more recently by the discovery of genes that encode fused remote homologs of the four eukaryotic histones in *Marseilleviridae*, a subfamily of giant viruses that infect amoebae. We demonstrate that viral doublet histones localize to the cytoplasmic viral factories after virus infection, and ultimately to mature virions. CryoEM structures of viral nucleosome-like particles show strong similarities to eukaryotic nucleosomes, despite the limited sequence identify. The unique connectors that link the histone chains contribute to the observed instability of viral nucleosomes, and some histone tails assume structural roles. Our results further expand the range of ‘organisms’ that have nucleosomes and suggest a specialized function of histones in the biology of these unusual viruses.

**One Sentence Summary:** Some large DNA viruses encode fused histone doublets that are targeted to viral factories and assemble into open nucleosome-like structures.

## Main Text

The organization of genomic DNA with histones into distinct complexes known as nucleosomes is a universal and highly conserved feature of all eukaryotes. The eukaryotic nucleosome core invariably comprises two copies each of the four unique histones H2A, H2B, H3 and H4. Each protein has a histone fold (HF) region that is structurally conserved between the four histones, as well as additional HF extensions and highly charged cationic tails that are unique to each (Luger and Richmond, 1998b). H2B-H2A and H4-H3 form obligate heterodimers which assemble into an octamer that wraps 147 base pairs (bp) of DNA to form nucleosomes (Luger et al., 1997).

The evolutionary origin of eukaryotic chromatin organization is widely thought to lie in the archaeal domain of life (Talbert et al., 2019). Archaeal histones are limited to the histone fold and are encoded by only one or a few closely related genes. They bind and bend DNA as homodimers or quasi-symmetric heterodimers using architectural principles also seen in eukaryotic histones but lack the ability to form defined particles and instead exist in ‘slinky-like’ assemblies that organize between 90 and ^~^ 600 bp of DNA (Bowerman et al., 2021; Mattiroli et al., 2017).

The genomes of some viruses (in particular nuclear DNA viruses and retroviruses that need to evade the host DNA damage recognition machinery) are organized into nucleosomes by appropriating eukaryotic histones and the host nucleosome assembly machinery during the latent and early lytic phase (e.g. (Oh et al., 2015), reviewed in (Lieberman, 2008)). To our knowledge, no histone proteins are found in the capsids of any of these viruses, nor do they encode viral histone homologues. In contrast, distinct histone-like proteins with homology to eukaryotic H2A, H2B, H3, and H4 have been identified in the genomes of some nucleo-cytoplasmic large DNA viruses (NCLDV). Several members of the *Marseilleviridae* isolated from the Amoeba *Acanthamoeba castellanii (A. castellanii)* encode homologues of the four histone proteins in native doublet form, where H4 is fused to H3, and H2B is fused to H2A (Fig. 1A) (Thomas et al., 2011). These proteins are present in Marseillevirus virions (Boyer et al., 2009; Fabre et al., 2017; Okamoto et al., 2018), where they might participate in the organization of the large (>300 kb) viral genomes. The arrangement of histones in doublets, the low level (<30 %) of sequence similarity with eukaryotic histones, and their high degree of conservation within the family of *Marseilleviridae* (Fig. 1A and Fig. S1A), suggests that they have evolved to fulfill viral functions other than (or in addition to) DNA compaction. This prompted us to investigate whether viral histones associate with the viral factory, whether they interact with DNA, and what types of structures they might form. We show that native histone doublets from Melbournevirus, a member of *Marseilleviridae* family of giant viruses, are specifically targeted to viral DNA and viral particles in Amoebae but are at most inefficiently incorporated into eukaryotic host chromatin. Using cryogenic electron microscopy (cryoEM), we demonstrate that viral histone doublets indeed interact with DNA to form nucleosome-like particles with distinct structural properties not seen in eukaryotes.

**Fig. 1.**
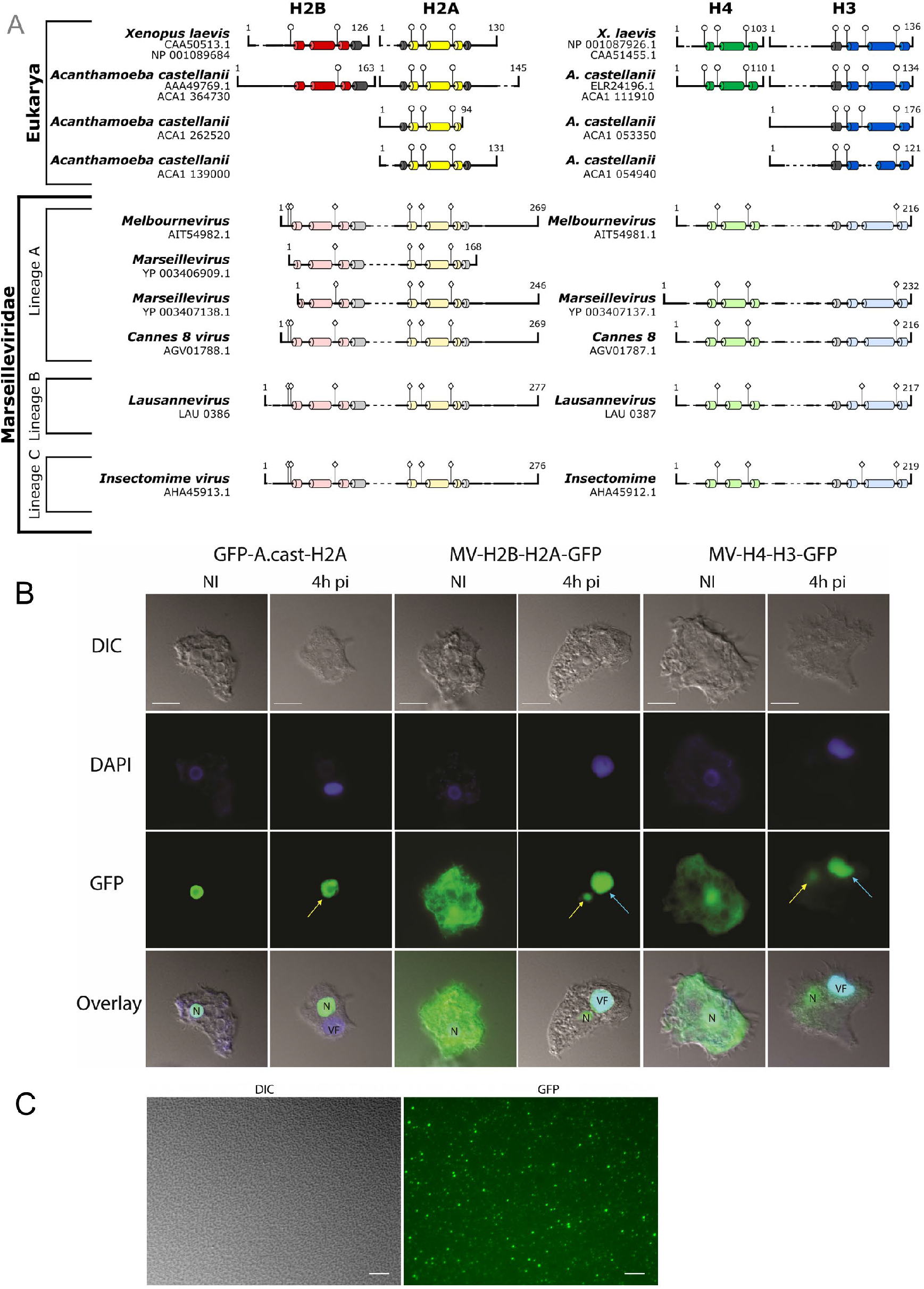
Melbournevirus histone doublets are re-localized to the viral factories (VF) and are loaded in the mature viral particles. (A) Histone dimer pairs (H2B, H2A and H4, H3) within Eukarya were aligned against the doublet *Marseilleviridae* histones using HHPRED’s multiple sequence alignment tool (ClustalΩ). Known α helices from the histone fold domain in Eukarya are dark colored tubes; H2B red, H2A yellow, H4 green, H3 blue, and additional helices in grey. Predicted α helices in MV-histones were generated using HHPRED’s Quick 2D prediction web server (shown in lighter coloration) within the *Marseilleviridae* histone doublets. Known R-T pairs and DNA binding residues are shown in Eukarya histones along with their conservation within *Marseilleviridae* histones; includes additional predicted DNA binding residues (positions demonstrated by lollipops). **(B)** Light microscopy fluorescence images (scale bar 10 μm) of *A. castellanii* cells transfected with GFP-*A. castellanii-H2A*, MV-H2B-H2A-GFP and MV-H3-H4-GFP, non-infected and infected with Melbournevirus at 4h pi. While GFP-*A. castellanii-H2A* concentrates only in the nucleus (N) of the non-infected cells, MV-H2B-H2A-GFP and MV-H3-H4-GFP are scattered in the entire cell (including the nucleus). Upon virus infection, GFP-*A. castellanii-H2A* remains in the nucleus (yellow arrows) while MV-H2B-H2A-GFP and MV-H3-H4-GFP re-localize into the viral factory (VF – cyan arrows). DAPI staining remains in the nucleus all along the infection but the intense fluorescence in the late VF hides the staining of the nucleus at 4h pi. **(C)**. Differential Interference Contrast (DIC) and GFP-fluorescence microscopy of Melbournevirus mature viral particles (scale bar 10 μm) produced in *A. castellanii* cells transfected with MV-H4-H3-GFP shows that MV-H4-H3-GFP is present in the mature viral particles.

### Melbournevirus histone doublets accumulate in virus factories and mature virions

Melbournevirus (MV) encodes three putative histone doublets (MEL_369, MEL_368, and MEL_149, here named MV-H2B-H2A, MV-H4-H3 and MV-miniH2B-H2A). Putative histone domains in each doublet are linked by a ^~^20 amino acid connector (Fig. 1A). Although these doublet histone proteins share less than 30 % amino acid sequence identity with eukaryotic histones, they are highly conserved within the group of *Marseilleviridae* (Fig. S1A). Secondary structure prediction suggests that viral histones form histone folds (α1-L1-α2-L2-α3) but have either longer (MV-H2B-H2A) or shorter (MV-miniH2B-H2A) H2A C-terminal tails than their eukaryotic counterparts (Fig. 1A). The H3 αN helix (which organizes the terminal turn of eukaryotic nucleosomal DNA) is predicted to be present in the viral H4-H3 doublets, while the sequence of the H2A docking domain (which tethers the H2A-H2B dimers to the (H3-H4)_2_ tetramer in eukaryotic nucleosomes) diverges from eukaryotic H2A (Fig. S1A). Many of the ‘signature’ amino acids that are important for DNA binding in eukaryotic histones are conserved in viral histones. These include arginine side chains that reach into the DNA minor groove and, in MV-H4-H3 interact with a threonine from the paired L1 loop (R-T pair) as well as many other basic amino acids (Fig. 1A, S1A).

To identify the localization of Melbournevirus histone doublets during viral assembly, we visualized transfected cells expressing fluorescently tagged MV-histone doublets as well as tagged Amoeba histones at different time points post infection (pi), and without virus infection. Amoeba GFP-H2A concentrated only in nuclei, where it presumably associates with genomic DNA to form nucleosomes, regardless of Melbournevirus infection (Fig. 1B, S1B). In contrast, unique and distinct patterns were observed for the transfected MV-histone doublets upon virus infection, particularly in the virus factory (VF; foci in the Amoeba cytoplasm where viral transcription, replication and assembly take place). MV-H2B-H2A-GFP and MV-H4-H3-GFP were initially dispersed throughout the whole cell including the nucleus, but their localization started to change between 1 and 2 hours post infection (Fig. S1C, D). Eventually, MV-H2B-H2A-GFP and MV-H4-H3-GFP accumulated and concentrated predominantly as foci in the viral factory (Fig. 1B and S1C, D; cyan arrows), although still present in nuclei at much lower concentrations (Fig. 1B, yellow arrows). Similar localization patterns were obtained with MV-miniH2B-H2A-GFP (Fig. S1E). Subsequently, fluorescently labeled MV-histone doublets were integrated into the mature virions (Fig. 1C). Altogether, the distinct localization pattern reveals specific targeting of Melbournevirus histone doublets to the viral DNA and the viral particle.

### Melbournevirus doublet histones form defined, unstable nucleosome-like particles

To determine what types of complexes MV-H2B-H2A, MV-miniH2B-H2A and MV-H4-H3 doublets form with DNA, we expressed, purified, and refolded the proteins from *E. coli* (Fig. S2A). When combined with DNA (‘601’ nucleosome positioning sequence) of varying lengths, and using the classic salt-gradient nucleosome reconstitution protocol (Dyer et al., 2004), MV-H2B-H2A and MV-H4-H3 form defined nucleosome-like particles (MV-NLP) that are stable at 37 °C, whereas individual histone doublets fail to form defined bands on DNA (Fig. 2A). The composition of the MV-NLP was confirmed by sucrose gradient fractionation (Fig. S2B). MV-histones also formed defined particles on a native 181 bp DNA fragment derived from the Melbournevirus genome (GC content = 45%). Irrespective of DNA sequence, and unlike *X. laevis* nucleosomes (eNuc), MV-NLPs dissociate upon heat treatment at 55 °C (Fig. S2C). MV-miniH2B-H2A alone binds DNA poorly, and no homogeneous MV-NLPs were formed with MV-H4-H3 and DNA (Fig. S2D). We therefore focused on MV-NLP with full-length MV-H2B-H2A and MV-H4-H3.

**Fig. 2.**
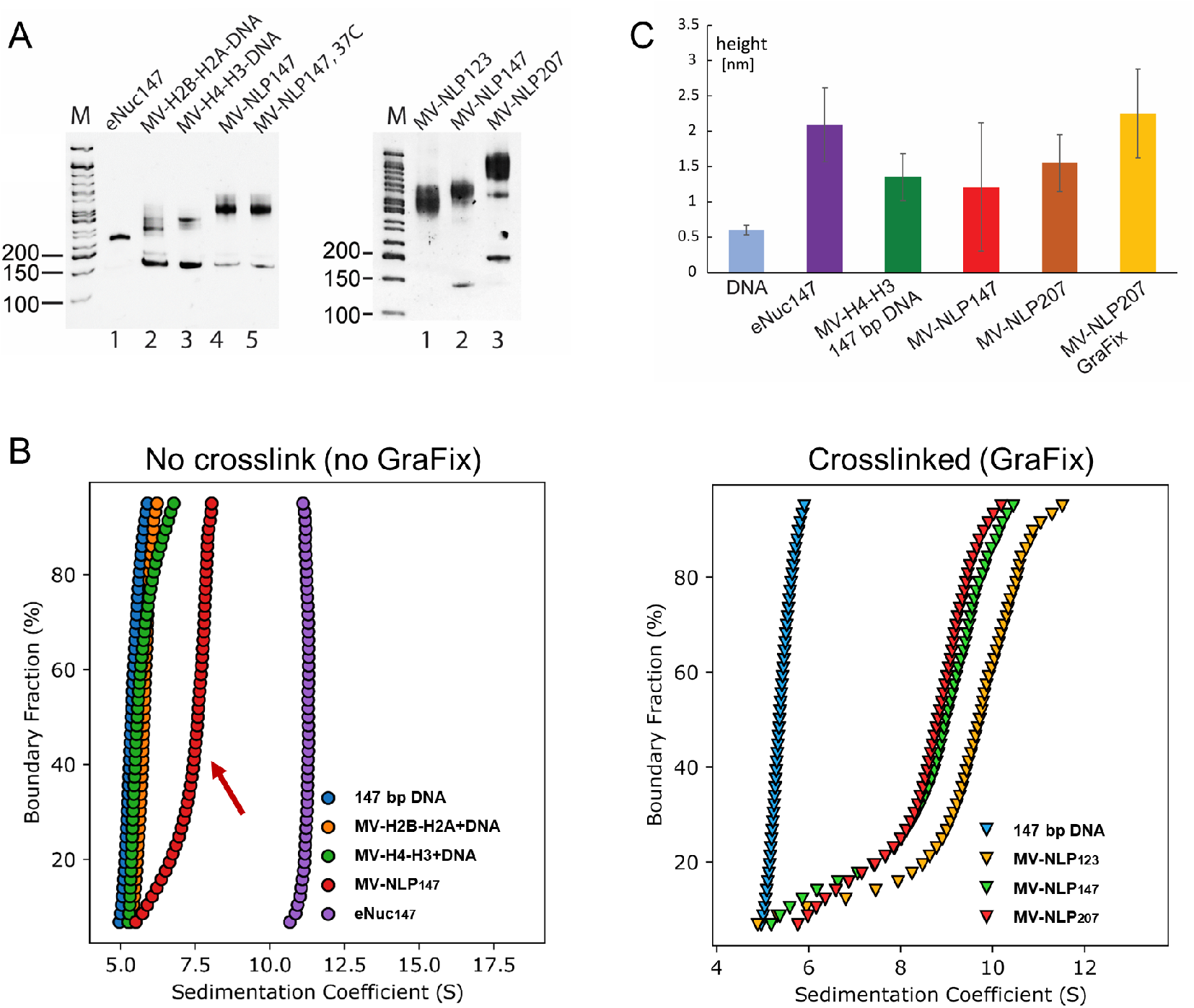
Histone doublets in *Marseilleviridae* form nucleosome-like particles (MV-NLP). **(A)** Native PAGE of reconstituted MV-NLPs with ‘601’ DNA of various lengths. Left panel: individual and combinations of histone doublets reconstituted onto 147 bp DNA; right panel: MV-H2B-H2B and MV-H4-H3 reconstituted onto 601 DNA of varying lengths. **(B)** Sedimentation Velocity Analytical Ultracentrifugation (SV-AUC) of MV-NLPs. Left: van Holde-Weischet plot of eukaryotic nucleosomes (eNuc), histone-DNA complexes with individual Melbournevirus histone doublets, and native MV-NLP with 147 bp DNA (no GraFix); right: van Holde-Weischet plot of crosslinked (GraFix-ed) MV-NLPs with 123, 147 or 207 bp DNA. **(C)** Height profile of MV-NLPs with 147 or 207 bp DNA, obtained by Atomic Force Microscopy (AFM). The average height profiles with standard deviations of the particles from each sample are shown; representative, original images are shown in supplementary data, statistics are shown in Table S1.

Sedimentation velocity analytical ultracentrifugation (SV-AUC) yields information on the size and shape distribution of macromolecular assemblies in solution. The diffusion-corrected sedimentation value of a particle is proportional to its mass, and inversely proportional to its viscous drag. The sedimentation behavior of MV-histone doublets assembled onto 147 bp DNA was compared to that of *X. laevis* histones reconstituted on the same DNA (eNuc_147_). The eNuc_147_ particle sediments at 11 S (Wang et al., 2018), while MV-NLP_147_ sediments at ^~^ 8 S, with a ‘tail’ towards lower S-values indicating dissociation of DNA at the relatively low (300 nM) concentrations used for AUC experiments (Fig. 2B). Particles reconstituted with individual MV-histone doublets sediment at ^~^5.5 S (Table 1). The AUC-derived molecular mass calculated for MV-NLP_147_ (Fig. 2B, red arrow) suggests that at least one copy of the H2B-H2A doublet has been lost due to the sample dilution in AUC (Table 1). To counteract the apparent dissociation of MV-NLPs, we employed Gradient Fixation (GraFix) with glutaraldehyde (Kastner et al., 2008). Histone doublets were efficiently crosslinked and the position of the main peak fraction in the gradient as well as migration on a native gel were unchanged upon crosslinking, suggesting little to no structural changes (Fig. S2E). Particles reconstituted on 123, 147, and 207 bp DNA fragments sedimented between 8.8 and 9.9 S after crosslinking (Fig. 2B, Table 1). The frictional ratio (f/f0) - a measure of particle ‘extension’, where larger values indicate increased drag, slowing sedimentation - of all MV-NLPs was higher than that of eNuc_147_, and f/f0 values increase in correlation with DNA length (Table 1). MV-NLP_123_ has the highest S-value among MV-NLP constructs, despite having the smallest mass, because of its significantly lower f/f0 value. This suggests that the additional DNA in MV-NLP_147_ and MV-NLP_207_ is extended.

**Table 1.** S values (S_(20,W)_), frictional ratios (f/f0) and calculated molecular weights (including confidence intervals) of histone-DNA complexes derived from SV-AUC.

|  | Sample | S(20,W) | f/f<sub>0</sub> | Molecular weight (kDa)<br>Exper./Theoretical |
| --- | --- | --- | --- | --- |
| No<br>GraFix | 147bp DNA | 5.10 (5.09, 5.10) | 3.09 (3.09, 3.09) | **117.13** (116.9, 117.3) / **97** |
|  | MV-H2B-H2A+DNA | 5.68 (4.61, 6.75) | 2.66 (2.51, 2.81) | **134.61** (89.83, 179.38) / **150** |
|  | MV-H4-H3+DNA | 5.28 (4.59, 5.96) | 2.79 (2.70, 2.87) | **120.93** (104.14, 137.71) / **146** |
|  | MV-NLP147 | 7.78 (7.74, 7.82) | 2.0 (1.96, 2.04) | **179.17** (173.55, 185.80) / **206** |
|  | eNuc147 | 11.16 (11.15, 11.16) | 1.54 (1.52, 1.56) | **218.5** (214.4, 222.5) / **210** |
| GraFix | MV-NLP123<br>(nucleosome-like) | 9.86 (7.99, 11.72) | 1.77 (1.52, 2.02) | **221.46** (170.93, 271.99) / **190** |
|  | MV-NLP147<br>(nucleosome-like) | 8.79 (7.30, 10.29) | 2.01 (1.86, 2.14) | **217.79** (140.57, 295.00) / **206** |
|  | MV-NLP207<br>(nucleosome-like) | 9.01 (7.48, 10.55) | 2.26 (2.00, 2.51) | **256.58** (194.54, 318.62) / **245** |

We visualized MV-NLPs by Atomic Force Microscopy (AFM). Particles reconstituted with either MV-H4-H3 alone or with the full complement of histone doublets were deposited onto mica surfaces and their height profiles were compared to eNuc_147_ (Fig. 2C, S2F, Table S1). The majority of eNuc_147_ survived the low concentration (2 nM) required for AFM imaging and presented the characteristic height profile of ^~^2.1 nm (White et al., 2016). In contrast, over 30% of the observed particles heights indicative of free DNA, indicating complete MV-NLP_207_ disassembly, and the remaining particles had lower height profiles (^~^1.3 nm) than the eNuc_147_ complex (Fig. 2C, S2F, Table S1). This reduced height suggests that MV-NLP_207_ transiently disassemble at the low concentration required for AFM. Crosslinking MV-NLP prevented disassembly and resulted in particles comparable in height to eNuc_147_.

### MV-NLPs resemble eukaryotic nucleosomes

Single particle electron cryo-microscopy (cyroEM) was used to visualize crosslinked MV-NLP_207_. GraFix was required as the majority of untreated MV-NLP dissociate during plunge freezing. Raw images and 2D classes show that MV-NLPs share many characteristic features of eukaryotic nucleosomes (Fig. S3A, B), with defined segments of extending linker DNA visible in many classes. After 3D classifications and refinement, we obtained a structure of the MV-NLP_207_ at 6.5 Å (Fig. S3C, D, Table S2). A second dataset obtained with GraFix-treated particles reconstituted onto 147 bp DNA (MV-NLP_147_) yielded an improved overall resolution of 4.2 Å (Fig. S4). In both structures, electron density for the DNA and histone helices are clearly distinguishable (Movie S1 and S2), which allowed the assignment of ^~^ 120 bp of bound DNA and all histone chains (Fig. 3A, Movie S3).

**Fig. 3.**
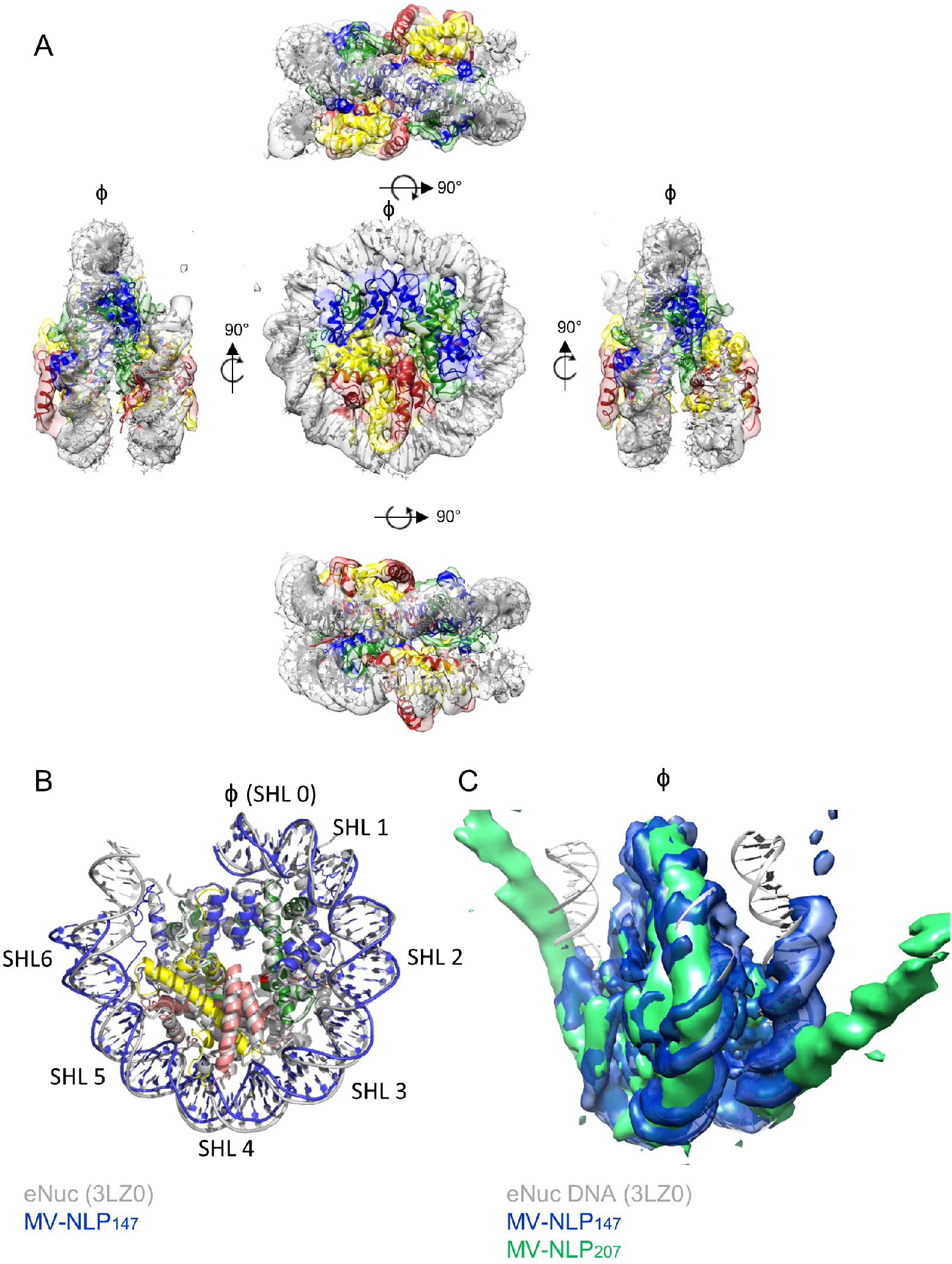
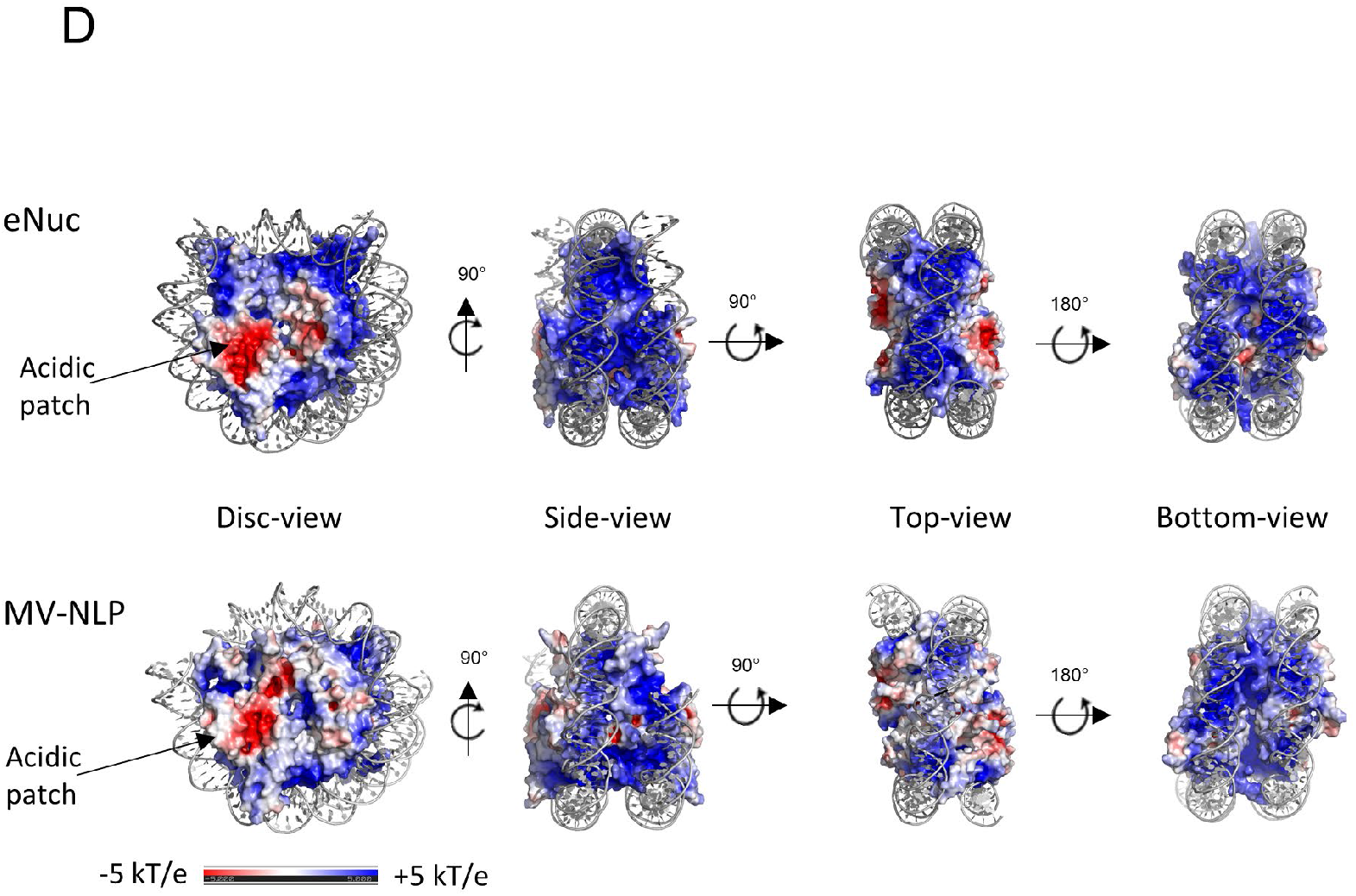
CryoEM reveals that MV-histone doublets form nucleosome-like structures with asymmetrically extending DNA. **(A)** Overview of MV-NLP_147_ and electron density. The equivalent regions of MV H3, H4, H2A, and H2B are shown in blue, green, yellow, and red, respectively. **(B)** Overlay of MV-NLP_147_ (blue) with eNuc (grey). Only 80 bp of DNA with associated histones are shown for clarity. Superhelix locations (SHL) are numbered from 0 to 6 starting from the nucleosome dyad (φ). (C) Comparison of the DNA path of eNuc_147_ (grey ribbon diagram), MV-NLP_147_ (blue electron density) and MV-NLP_207_ (green electron density). (D) Charged surface representation of the histones for MV-NLP_147_ and eNuc_147_. Coordinates for eNuc_147_ taken from 3LZ0.

The overall dimensions of the DNA superhelix, the path described by the DNA around the histone core, and the overall layout of histone fold helices are similar between MV-NLPs and eukaryotic nucleosomes (Fig. 3A, B; Table S3). However, DNA wrapping is incomplete and asymmetric in both MV-NLPs, as only ^~^ 120 bp are wrapped in most particles. This is more pronounced in MV-NLP_207_, where well-defined electron density describes a straight path for DNA extending away from the histone core at superhelical location (SHL) 4.5, whereas the DNA extending at the other side (from SHL 5.5) appears to be more disordered (Fig. 3C, Movie S1). In MV-NLP_147_ complex slightly more DNA is bound (up to SHL 5.5 on both sides), and only one arm of extending DNA has defined density (Fig. 3C, Movie S2). This asymmetry is likely a consequence of the asymmetric 601 DNA sequence (Chen et al., 2017; Chua et al., 2012).

Initial models of MV histone doublets were generated though homology modeling (Waterhouse et al., 2018), where the ^~^28 and ^~^20 amino acid connectors in H4-H3 and H2B-H2A were constructed through *de novo* methods (Webb and Sali, 2016). These were docked into the MV-NLP_147_ density, with good agreement with nucleosome-like configurations (Fig. 3A, B; correlation coefficient between model and experimental density 0.766).

Overall, the MV-histone core is less positively charged than the eukaryotic histone octamer (pI of ^~^9.5 vs. ^~^11.0; Fig. 3D). While basic amino acids describe a distinct path for the DNA in both histone cores, and many amino acid side chains that engage DNA are conserved between viral and eukaryotic histones (Fig. 1A, S1A), positive charges along the helical path are less pronounced in MV-histones, in particular in the region formed by the H3-H3’ four-helix bundle (‘dyad’). This contributes to the reduced stability of the MV-NLP compared to eNuc. Additionally, the acidic patch, the primary point of interactions between eukaryotic nucleosomes and nuclear proteins (Peng et al., 2020), differs in size and charge, due to fewer acidic residues in MV-H2A (Fig. 3D).

The two MV-H4-H3 doublets superimpose onto the eukaryotic (H4-H3)_2_ tetramer with a backbone rmsd of ^~^2 Å (Fig. 4A; Table S3). The main chain arrangements of the MV-H3-H3’ four helix bundles at the nucleosomal dyad are similar as in eNuc_147_, but the interface lacks the hallmark histidine – cysteine configuration. Each MV-H2B-H2A doublet interfaces with the H4 portion of a MV-H4-H3 doublet through a four-helix bundle of similar architecture as the H4-H2B interface in eNuc_147_. However, the MV-H2B α2 helix is one turn shorter than the eukaryotic sequence (due to a conserved proline at position 83), and the tyrosines that form π-stacking interactions in eNuc systems are consistently absent in both MV-H4 and MV-H2B (Fig. 4B). Additionally, while the main chains of the two H2A L1 loops (^38^NYAE^41^) are close to one another in eNuc_147_ and form direct interactions through their side chains, the distance between L1 loops in MV-NLP_147_ (^153^GGCS^156^; positions 214-217 in Fig. S1A) is increased and shows no direct L1-L1 interactions. L1-L1 separation is accompanied by a similar increase in the distance between H2B-H2A centers (36.3 Å in eNuc_147_ vs 38.6 Å in MV-NLP_147_; Table S3). In addition to being further apart in the MV-NLP_147_ complex, the H2B-H2A histone folds reorient away from the nucleosome core (Fig. 4C, Fig. S5A, Table S3). Conversely, the H2A docking domains of MV-H2A are angled inward, toward the dyad axis, in an altogether different arrangement than what is seen in eNuc_147_ (Fig. 4C, Table S3; 6-8 Å backbone RMSD for chains C and G, respectively). Together, these structural changes may contribute to the reduced stability, and increased propensity to form sub-NLPs (hexasomes and tetrasomes), of MV-NLPs in comparison to the eNuc complex.

**Fig. 4.**
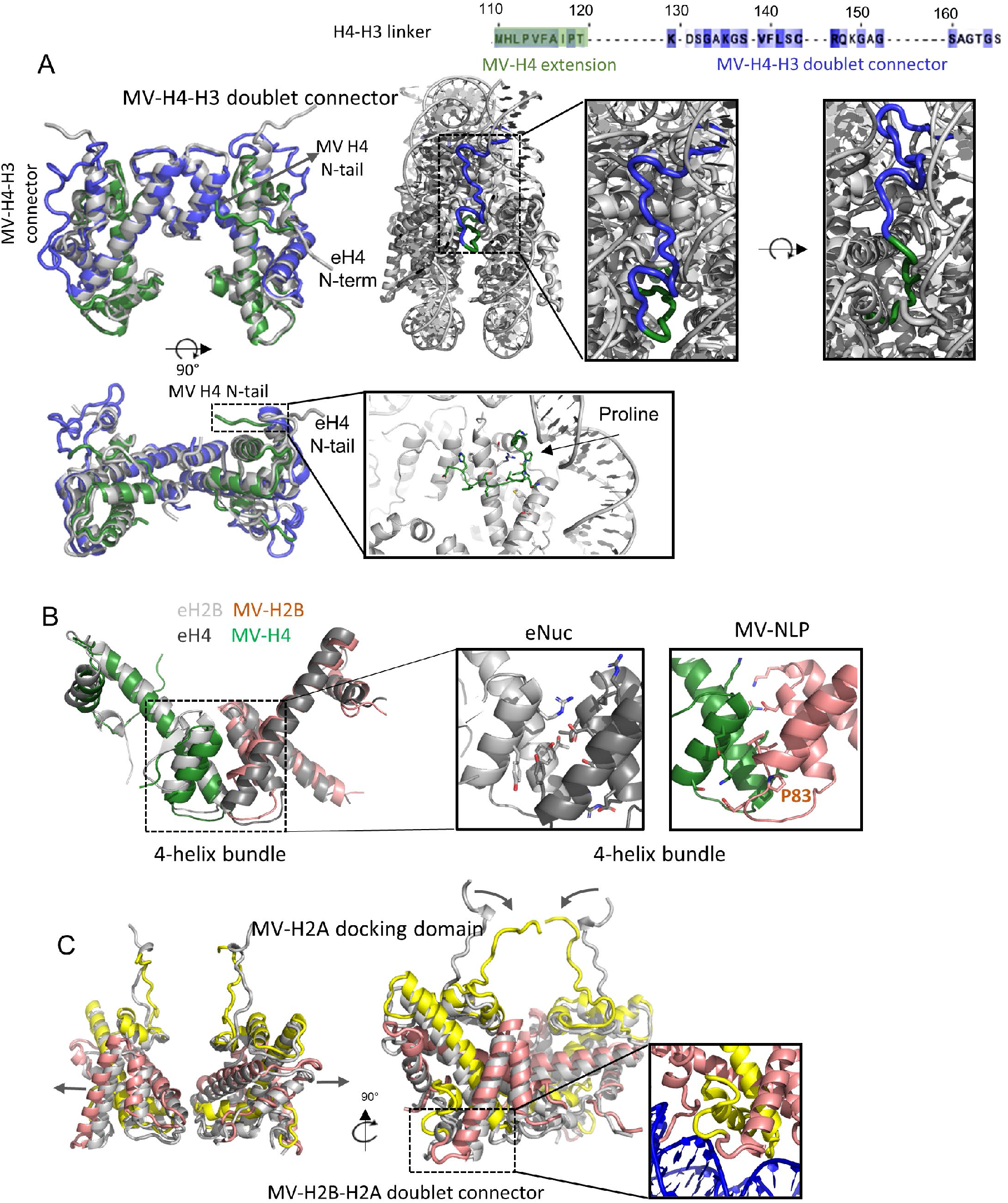
Comparison of MV-NLP and eNuc histone structures. (A) Superposition of two MV-H4-H3 doublets (green and blue) with the eukaryotic (H3-H4)_2_ tetramer in grey (left), and close-up of MV-H4-H3 doublet connector (right); Interactions of the MV-H4 N-terminal tail with the nucleosome are also shown. (B) A comparison of the four-helix bundle structure formed by H4 and H2B. (C) Superposition of two MV-H2B-H2A doublets (in red and yellow) with eukaryotic H2A-H2B dimers in grey (left), and close-up of the MV-H2B-H2A doublet connector and docking domain. Additional possible configurations for both connectors are shown in Fig. S5.

The connector linking the C-terminus of MV-H2B with the N-terminus of MV-H2A is easily accommodated as these are near each other. A variety of favorable conformations are predicted through *de novo* modeling, with several different arrangements in agreement with observed weak density in MV-NLP_147_ (Fig. 4C inset, S5B). A conserved arginine is pointing into the minor groove, and main-chain phosphate interactions further hold the loop in place, echoing the contributions of the H2A N-terminal domain in eNuc. Connecting the MV-H4 C-terminus with the N-terminal tail of MV-H3 is conceptually more difficult. In eukaryotic H4, the eight most C-terminal amino acids are engaged in contacts with H2A and H2B in the interior of the histone core. Our models show that the equivalent region in MV-H4 maintains docking domain interactions with MV-H2A. Physical constraints dictate that the missing linker segment must continue toward the DNA superhelix and project upward in the direction of MV-H3 αN, with which it connects (Fig. 4A). Electron density attributed to the connector is visible between the DNA gyres on both sides, and in both structures (MV-NLP_207_ and MV-NLP_147_). Although not sufficiently defined to allow modeling through real-space refinement, *de novo* generation of linker structures provided multiple feasible conformations (Fig. S5C), and molecular dynamics flexible-fitting simulations confirmed that these orientations are physically relevant (Fig. 4A). A stretch of 10 predominantly hydrophobic amino acids, packs against the region connecting MV-H3αN with α3 to form a hydrophobic core (Fig. 4A, Fig. S5C), and the connector is also positioned to engage in main-chain contacts with the DNA backbone at SHL +/- 1.5. Mostly flexible and small amino acids (GAGSAGTGS) form the turn towards MV-H3 αN (Fig. 4A). As such, the MV-H4-H3 connector (which is highly conserved among *Marseilleviridae* doublets) can be accommodated within the structural framework of a canonical nucleosome, although its presence likely contributes to the unwrapping of the terminal 10 bp of DNA observed in both MV-NLP densities.

The other MV-histone tails that are not engaged in doublet formation also differ from the histone tails in eNuc. The C-terminal tail of MV-H2A is significantly longer than that of eukaryotic H2A (Fig. S1A). The MV-H4 N-terminal domain (equivalent to the first 26 amino acids in eH4) does not extend over the DNA but takes the opposite direction to interact with MV-H4 α2 and α3 (Fig. 4A). This density is as well-defined as that of the main histone fold helices, suggesting that this domain may consistently interact with its own histone core rather than with DNA or the acidic patch in neighboring particles, as observed for the eukaryotic H4 tail (Dorigo et al., 2003; Morrison et al., 2018). The amino acid residues in this region are highly conserved among *Marseilleviridae* and differ from the eH4 tail (Fig. S1A). A conserved proline (P34) at the tip of MV-H4 α1 might be responsible for the redirection (Fig. 4A).

As is the case with the eH4 tail, the first 20 amino acids of the MV-H4 tail are too disordered to be observed in the electron density map.

## Discussion

The discovery that the genomes of some giant viruses encode histone-like proteins in which the dimerization partners are fused into a single chain was surprising (Thomas et al., 2011). Our results that these MV-histone doublets associate with the viral factory and indeed with the mature virus, and that they form nucleosome-like particles with unique properties, supports the idea that they are involved in viral genome organization. Instead of organizing DNA into nucleoprotein ‘core complexes’ inside the viral capsid, *Marseilleviridae* utilize their own specialized histone doublets to form nucleosomes and possibly more compact chromatin within the virus particle. In the context of the ongoing debate regarding the role of viruses in the emergence of the eukaryotic nucleus, our findings also provide a new dimension to the diverse role of histones in genome organization of non-eukaryotic entities (Erives, 2017).

*Marseilleviridae* exhibit intermediate dependency on their host as they develop their viral factory in the cytoplasm but transiently recruit host nuclear proteins to the viral factory to transcribe early genes, until the virally encoded RNA polymerase has been synthesized (Fabre et al., 2017). This is achieved through a reorganized leaky nucleus during the early phase. Amoeba histones are exclusively localized in the host nucleus whether the cells are infected or not, whereas viral histones are present in the nucleus and cytoplasm in uninfected cells but move into viral factories upon infection to integrate into mature virions. This suggests that Amoeba histones remain bound to cellular DNA, while the viral histones that made it into the nucleus apparently do not interact with Amoeba genomic DNA and can leave the nucleus to associate with the viral factory. As such, Amoeba histones are likely targeted to the nucleus and assembled into chromatin by histone chaperones and transporters that mostly avoid viral histones. As is the case for the other nuclear proteins transiently leaving the cell nucleus upon *Marseilleviridae* infection, it remains to be seen whether viral histones are actively or passively recruited to the VF.

Many ‘nucleosome signature features’ are maintained between archaeal, eukaryotic and now viral nucleosomes, such as the overall geometry and arrangement of the DNA superhelix, achieved by interactions between the DNA minor groove backbone and the main chain of antiparallel ‘L1L2’ loops (from H4-H3 and H2B-H2A pairs), and the utilization of positive α helix dipole moments to bind DNA (Luger and Richmond, 1998a). Several of the ‘sprocket arginines’ (Hodges et al., 2015) and their pairing with a threonine from the dimerization partner, as well as the highly conserved R-D clamps stabilizing the histone fold, are conserved between eukaryotic, archaeal, and viral histones. However, MV-doublet histones stably bind only ^~^ 120 bp of DNA as opposed to the 147 bp in eukaryotic nucleosomes. In our cryoEM structures, the unbound DNA is characterized by distinct density indicating that it is in a defined orientation. Unique to MV-NLP is also the packing of the H4 N-terminal tail onto the surface of the histone core, the overall less positive charge of the histone core surface including the acidic patch, and, most notably, the linking of MV-H4-H3 and MV-H2B-H2A to form doublet histones. Fused histone genes are also observed in some archaea (Talbert et al., 2019), but to date not in any eukaryotic genome. The amino acid linkers connecting the histone moieties to form the doublets, which are conserved in sequence across *Marseilleviridae*, can be accommodated in the structure. However, the MV-H4-H3 connector (which is conserved in length and amino acid sequence between members of the *Marseilleviridae*) likely contributes to destabilizing the last turn of DNA. Overall, MV-NLPs are significantly destabilized compared to eNuc and this seems to be an intrinsic property related to their virus-specific function.

A common trait of DNA viruses is that they must actively package their DNA into capsids to protect it outside the host cell environment. On the other hand, they also need to make their genomes accessible to the transcription machinery immediately upon infection. Previous analyses of the infectious cycle of Marseilleviruses revealed that the capsids appear to dissolve around 2h post infection, leaving a spherical electron dense core in the cytoplasm (Fabre et al., 2017). Our study suggests that *Marseilleviridae* assemble virally encoded histones into nucleosomes to condense and protect their genome, to allow it to fit into the ^~^200 nm diameter icosahedron, also explaining the higher than expected electron density of the spherical core (Fabre et al., 2017). Arguably, viral nucleosomes must be metastable to make the genome, once transferred into the cytoplasm, accessible to transcription by the host RNA polymerase recruited to the viral factory. This might explain the requirement for virus-encoded histones, as ATP-dependent remodelers utilized by eukaryotes to facilitate transcription through chromatin are localized mostly in the nucleus. The regions in MV-histones that convey this metastability (such as the H4-H3 connector, and the distinct makeup of the four-helix bundle regions) are highly conserved among *Marseilleviridae*. Why these histones exist as doublets, what determines their specificity for the viral genome, and which (if any) assembly factors they rely on for their association with the viral DNA are intriguing open questions that warrant further research.

## Acknowledgments

We thank Garry Morgan (CU Boulder); Rui Yan and Ziheng Yu (Janelia Research Campus cryoEM facility) for help with data collection.

## Author contributions

YL: conceptualization, methodology, validation, formal analysis, investigation, data curation, writing (Original draft, review and editing), visualization, supervision
CMT: Methodology; Validation; Formal Analysis; Investigation; Data curation; Writing (Review & Editing); Visualization
NP: investigation (in vivo data), data analysis
SJ: investigation (in vivo data), data analysis
KZ: Methodology; Validation; Formal Analysis; Data curation; Writing; Visualization
SB: Methodology; Software; Formal Analysis; Investigation; Data Curation; Writing - Review & Editing; Supervision; Funding Acquisition
AW: investigation, formal analysis, data curation
GE: investigation, formal analysis, data curation
CA: Conceptualization, formal analysis, writing (discussion and editing), Supervision; Project Administration; Funding Acquisition
KL: Conceptualization; Writing (Original Draft; Review & Editing); Supervision; Project Administration; Funding Acquisition

## Data and materials availability

All original data will be made accessible. Coordinates and xxx have been deposited in the appropriate databases (PDB and EMPIAR, access codes XXX and XXX). All other data is available in the main text or supplementary materials.

## Supplementary Materials

Materials and Methods

Figs. S1 to S6

Tables S1 to S3

Captions for Movies S1 to S3

Movies S1 to S3

## References

Bowerman, S., Wereszczynski, J., and Luger, K. (2021). Archaeal chromatin ‘slinkies’ are inherently dynamic complexes with deflected DNA wrapping pathways. Biorxiv.

Boyer, M., Yutin, N., Pagnier, I., Barrassi, L., Fournous, G., Espinosa, L., Robert, C., Azza, S., Sun, S., Rossmann, M.G., et al. (2009). Giant Marseillevirus highlights the role of amoebae as a melting pot in emergence of chimeric microorganisms. Proc Natl Acad Sci U S A 106, 21848–21853.

Chen, Y., Tokuda, J.M., Topping, T., Meisburger, S.P., Pabit, S.A., Gloss, L.M., and Pollack, L. (2017). Asymmetric unwrapping of nucleosomal DNA propagates asymmetric opening and dissociation of the histone core. Proc Natl Acad Sci U S A 114, 334–339.

Chua, E.Y., Vasudevan, D., Davey, G.E., Wu, B., and Davey, C.A. (2012). The mechanics behind DNA sequence-dependent properties of the nucleosome. Nucleic Acids Res 40, 6338–6352.

Dorigo, B., Schalch, T., Bystricky, K., and Richmond, T.J. (2003). Chromatin fiber folding: requirement for the histone H4 N-terminal tail. J Mol Biol 327, 85–96.

Dyer, P.N., Edayathumangalam, R.S., White, C.L., Bao, Y., Chakravarthy, S., Muthurajan, U.M., and Luger, K. (2004). Reconstitution of nucleosome core particles from recombinant histones and DNA. Methods Enzymol 375, 23–44.

Erives, A.J. (2017). Phylogenetic analysis of the core histone doublet and DNA topo II genes of Marseilleviridae: evidence of proto-eukaryotic provenance. Epigenetics Chromatin 10, 55.

Fabre, E., Jeudy, S., Santini, S., Legendre, M., Trauchessec, M., Coute, Y., Claverie, J.M., and Abergel, C. (2017). Noumeavirus replication relies on a transient remote control of the host nucleus. Nat Commun 8, 15087.

Hodges, A.J., Gallegos, I.J., Laughery, M.F., Meas, R., Tran, L., and Wyrick, J.J. (2015). Histone Sprocket Arginine Residues Are Important for Gene Expression, DNA Repair, and Cell Viability in Saccharomyces cerevisiae. Genetics 200, 795–806.

Kastner, B., Fischer, N., Golas, M.M., Sander, B., Dube, P., Boehringer, D., Hartmuth, K., Deckert, J., Hauer, F., Wolf, E., et al. (2008). GraFix: sample preparation for single-particle electron cryomicroscopy. Nat Methods 5, 53–55.

Lieberman, P.M. (2008). Chromatin organization and virus gene expression. J Cell Physiol 216, 295–302.

Luger, K., Mader, A.W., Richmond, R.K., Sargent, D.F., and Richmond, T.J. (1997). Crystal structure of the nucleosome core particle at 2.8 A resolution. Nature 389, 251–260.

Luger, K., and Richmond, T.J. (1998a). DNA binding within the nucleosome core. Current Opinion in Structural Biology 8, 33–40.

Luger, K., and Richmond, T.J. (1998b). The histone tails of the nucleosome. Curr Opin Genet Dev 8, 140–146.

Mattiroli, F., Bhattacharyya, S., Dyer, P.N., White, A.E., Sandman, K., Burkhart, B.W., Byrne, K.R., Lee, T., Ahn, N.G., Santangelo, T.J., et al. (2017). Structure of histone-based chromatin in Archaea. Science 357, 609–612.

Morrison, E.A., Bowerman, S., Sylvers, K.L., Wereszczynski, J., and Musselman, C.A. (2018). The conformation of the histone H3 tail inhibits association of the BPTF PHD finger with the nucleosome. Elife 7.

Oh, J., Sanders, I.F., Chen, E.Z., Li, H., Tobias, J.W., Isett, R.B., Penubarthi, S., Sun, H., Baldwin, D.A., and Fraser, N.W. (2015). Genome wide nucleosome mapping for HSV-1 shows nucleosomes are deposited at preferred positions during lytic infection. PLoS One 10, e0117471.

Okamoto, K., Miyazaki, N., Reddy, H.K.N., Hantke, M.F., Maia, F., Larsson, D.S.D., Abergel, C., Claverie, J.M., Hajdu, J., Murata, K., et al. (2018). Cryo-EM structure of a Marseilleviridae virus particle reveals a large internal microassembly. Virology 516, 239–245.

Peng, Y., Markov, Y., Goncearenco, A., Landsman, D., and Panchenko, A.R. (2020). Human Histone Interaction Networks: An Old Concept, New Trends. J Mol Biol.

Talbert, P.B., Meers, M.P., and Henikoff, S. (2019). Old cogs, new tricks: the evolution of gene expression in a chromatin context. Nat Rev Genet.

Thomas, V., Bertelli, C., Collyn, F., Casson, N., Telenti, A., Goesmann, A., Croxatto, A., and Greub, G. (2011). Lausannevirus, a giant amoebal virus encoding histone doublets. Environ Microbiol 13, 1454–1466.

Wang, T., Liu, Y., Edwards, G., Krzizike, D., Scherman, H., and Luger, K. (2018). The histone chaperone FACT modulates nucleosome structure by tethering its components. Life Sci Alliance 1, e201800107.

Waterhouse, A., Bertoni, M., Bienert, S., Studer, G., Tauriello, G., Gumienny, R., Heer, F.T., de Beer, T.A.P., Rempfer, C., Bordoli, L., et al. (2018). SWISS-MODEL: homology modelling of protein structures and complexes. Nucleic Acids Res 46, W296–W303.

Webb, B., and Sali, A. (2016). Comparative Protein Structure Modeling Using MODELLER. Curr Protoc Bioinformatics 54, 5 6 1–5 6 37.

White, A.E., Hieb, A.R., and Luger, K. (2016). A quantitative investigation of linker histone interactions with nucleosomes and chromatin. Sci Rep 6, 19122.

