## supplementary materials for "Melbournevirus-encoded histone doublets are recruited to virus particles and form destabilized nucleosome-like structures"

**This PDF file includes:**

Materials and Methods
Figs. S1 to S6
Tables S1 to S3
Captions for Movies S1 to S3

**Other Supplementary Materials for this manuscript include the following:**

Movies S1 to S3

#### Materials and Methods

##### Histone sequence alignment and secondary structure prediction

Predicted *Marseilleviridae* histone-like proteins were aligned with eukaryotic histone proteins with HHpred's Multiple Alignment using Fast Fourier Transform (MAFFT) with a 1.53 gap open penalty. Lineages A, B, and C of the *Marseilleviridae* family were included, lineage D was excluded due to sequence similarity with lineage C (~90%) and lineage E was excluded due to the lack of annotated histone-like proteins. Using the MAFFT alignment, protein secondary structures were predicted using HHpred's Quick 2D structural prediction webserver to demonstrate structural conservation of the histone fold domain between *Eukarya* and *Marseilleviridae* (*1*).

##### Fluorescence localization of Melbournevirus histones in infected *A. castellanii* cells

The three Melbournevirus (AIT54904) histones encoding genes Mel-368 (MV-H4-H3), Mel-369 (MV-H2B-H2A) and Mel-149 (MV-miniH2B-H2A) were amplified by PCR from genomic DNA and cloned into an in-house modified plasmid derived from the pGAPDH-GFP amoebal expression plasmid (*2*). Briefly, the GAPDH promoter was replaced by the stronger EF1 gene promoter and the genes were inserted using the NdeI restriction site to yield C-terminally GFP-tagged proteins. The *A. castellanii* histone H2A (2) gene (ACA1_364730A) was amplified from *Acanthamoeba castellanii* (Douglas) Neff (ATCC 30010TM) genomic DNA and cloned into the same plasmid engineered to yield a N-terminally GFP-tagged protein.

*Acanthamoeba castellanii* cells were transfected with 6 µg of each plasmid using Superfect (Qiagen). Selection of transformed cells was initially performed at 30 µg/mL Neomycin and increased up to 100 µg/mL within a couple of weeks. Transfected *A. castellanii* cells were grown on poly-L-lysine coating coverslips in a 12-well plate and infected with Melbournevirus at a MOI of 50 except for the negative control. At 30 min, 1h, 2h, 3h and 4h post-infection (pi), cells were fixed with PBS containing 3.7% formaldehyde for 20 min at room temperature. After one wash with PBS buffer, coverslips were mounted on a glass slide with 4 µl of VECTASHIELD mounting medium with DAPI and the fluorescence was observed using a Zeiss Axio Observer Z1 inverted microscope using a 63x objective lens associated with a 1.6x Optovar for DIC, DAPI or GFP fluorescence recording.

##### Observation of Melbournevirus fluorescent mature viral particles

*Acanthamoeba castellanii* cells transfected with MV-H4-H3-GFP were grown in two T175 flasks (about 100 000 cells/$cm^2$). Cells were infected with Melbournevirus at MOI 1. After cells lysis, the cultures were recovered and centrifuged for 10 min at 1 000 x g to remove the cell debris. The supernatants were filtered through 0.45 µm filters and centrifuged for 1h at 10,000 x g to pellet the viral particles. The viral pellet was resuspended in 2ml of PBS and filtered through a 0.22 µm filter. Fluorescence was observed using a Zeiss Axio Observer Z1 inverted microscope using a 63x objective lens associated with a 1.6x Optovar.

##### Melbournevirus histone doublet purification

Melbournevirus ORF 369 (H2B-H2A), ORF 368 (H4-H3) and ORF 149 (miniH2B-H2A) were cloned into pET-28 plasmid for expression and purification from *E.coli,* using protocols adapted from eukaryotic histones (*3*). 6L of *E. coli* cells were re-suspended and lysed with 80 mL of guanidinium lysis buffer (6 M Guanidinium HCl, 20 mM sodium phosphate pH 6.8 and 2 M NaCl) and sonicated for 30 seconds at 50% strength for 4 times. Lysate was spun at 16,000 rpm for 25 minutes, and the supernatant was incubated with Nickel NTA Agarose Beads (GoldBio) for approx. 45 minutes at RT. The nickel beads were pelleted, and the supernatant was discarded. The beads were resuspended in 8 M Urea lysis buffer (8 M Urea, 20 mM sodium phosphate pH 6.8, and 2 M NaCl) and incubated for approximately 1 hour at room temperature.

Proteins were eluted using histone elution buffer (8 M Urea, 20 mM sodium phosphate pH 6.8, 2 M NaCl, and 500 mM Imidazole). For protein refolding, histones were dialyzed against 1 M Urea Buffer (1 M urea, 1 mM EDTA, 20 mM Tris-Cl pH 7.5 and 2 M NaCl) for 6 hrs, followed by overnight dialysis against the same buffer without urea (1 mM EDTA, 20 mM Tris-Cl pH 7.5 and 2 M NaCl). Precipitated protein was removed by centrifugation, and the supernatant was concentrated and run over a size-exclusion column S200 equilibrated in 1 mM EDTA, 20 mM Tris-Cl pH 7.5 and 2 M NaCl. Samples were stored in 20% glycerol at -80°C.

**Marseillevirus histone-DNA complex (nucleosome) reconstitution**

Melbournevirus H2B-H2A, H4-H3 and DNA of the specified lengths were mixed at a ratio of DNA to MV-H4-H3 to MV-H2B-H2A = 1.0 : 2.4 : 2.4. Widom 601 DNA at 147 bp and 207bp, and 181 bp of a conserved Melbournevirus native DNA fragment selected from its genome (GC cotent = 45%) were used. Viral NLPs and *X.laevis* nucleosomes were both reconstituted by gradient dialysis (*3*). Reconstituted products were analyzed by 5 % native-PAGE.

**Sucrose gradient sedimentation and Gradient Fixiation (GraFix) crosslinking**

To form the sucrose gradient, 6 ml top solution (50 mM NaCl, 20 mM HEPES, 1 mM EDTA, pH 7.5 and 5% sucrose) was added to a tube (Beckman, 331372). Then 6 ml bottom solution (50 mM NaCl, 20 mM HEPES, 1 mM EDTA, pH 7.5 and 40% sucrose) was slowly added to the bottom until it reached the halfway mark. The tubes were placed on a gradient maker (BioComp Gradient Master) to form a continuous gradient. 200 µl of sample at 4 µM was loaded on top and spun at 4 °C for 18 h at 30,000 r.p.m. (Beckman, Rotor SW-41Ti). Fractioned samples were dialyzed against 50 mM NaCl, 20 mM Tris-Cl, 1 mM EDTA, pH 7.5 and 1 mM DTT to remove the sucrose.

For GraFix, the same procedure was applied, except that 10% to 30% of glycerol was used instead of sucrose. In addition, 0.15% of glutaraldehyde was added to the bottom solution to form the continuous density and crosslinker gradient. After GRAFIX, the samples were eluted and dialyzed against 50 mM NaCl, 20 mM Tris-Cl, 1 mM EDTA, pH 7.5 and 1 mM DTT to quench the crosslinking reaction and remove glycerol.

**Sedimentation Velocity Analytical Ultracentrifugation (SV-AUC)**

SV-AUC with absorbance optics ($\lambda$ = 260 nm) was used to evaluate the homogeneity of the complexes in solution and to infer their molecular size and shape. Samples (composition as indicated) at 300 nM were spun at 30–35,000 rpm at 20°C in a Beckman XL-A ultracentrifuge in 50 mM NaCl, 20 mM Tris-Cl, 1 mM EDTA, pH 7.5 and 1 mM DTT, using an An60Ti rotor. Partial specific volumes of samples were estimated using UltraScan III version 4.0; this program was also used for all data analysis (*4, 5*).

Time-invariant and radial-invariant noise contributions were subtracted from the experimental sedimentation velocity data by 2-dimensional Spectrum Analysis (2DSA), followed by refinement using the Genetic Algorithm-Monte Carlo (GA-MC) approach (*6*). Sedimentation coefficients and molecular weights in Table 1 were extracted from resulting GA-MC models. Modeling calculations were performed on the UltraScan LIMS clusters at the Bioinformatics Core Facility at University of Texas Health Science Center at San Antonio and Lonestar cluster at Texas Advanced Computing Center. Integral sedimentation coefficient distributions (G(s)) were obtained from noise-corrected experimental data with the Enhanced van Holde-Weischet Analysis method. G(s) plot figures were created using python (*7*) (https://github.com/Luger-Lab/AUC-analysis).

.

**Atomic Force Microscopy (AFM)**

All samples were imaged in air on JPK/Bruker NanoWizard Sa with TAP300-Gold (Ted Pella) cantilevers using fast scan glass block. Images were usually scanned at 1.5 x 1.5 µm at 1- 3 hertz. All buffers were filtered through 0.02 µM filters (Anotop) before use. Mica was freshly cleaved, treated with APTES for 30 min, rinsed with water and then dried with nitrogen gas utilizing a 0.22 µm PES filter. Samples were serially diluted in TE (20 mM Tris-HCl pH 7.5, 1 mM EDTA) to a final concentration of 2 nM. Within 2 minutes of final dilution, 30 µl sample was placed on the ATPES mica slide for 2 minutes, rinsed with 1.0 ml of water and dried with filtered $N_2$ gas. Images were analyzed using JPK SPM software V 6.1.158. Each 1.5 x 1.5 µm image was divided into 4 quadrants and digitally zoomed to obtain clearer visuals of separated particles. Lines were drawn and dragged through each particle with the greatest height of each recorded. Analysis of heights were completed in either MS Excel and/or GraphPad Prism.

**Single particle cryo Electron Microscopy (cryo EM) and data processing**

Complexes of MV-NLPs from GraFix were concentrated to 2-3 µM using Amicon Ultra-4 centrifugal filters (Ultracel 50K, Millipore). C-Flat 1.2/1.3 (Au) grids were glow discharged (EMItec, Lohmar, DE) at 40 mA for 30 s. 4 µl sample was applied onto the grid before manual plunge freezing into ethane. Images for MV-NLP with 147 bp "601" DNA were acquired at nominal magnification of 64000x on a FEI Titan Krios (300 kV), equipped with a Gatan K3 Summit direct detector. Pixel size was 1.065 Å. The movies were captured in super resolution mode with electron dose rate at 10.5 electrons per pixel per second for 5.4 s and 0.108 s per frame. Defocus range was −0.8 to −2.0 µm. Images for MV-NLP with 207 bp "601" DNA were acquired at nominal magnification of 29000x on a FEI Tecani F20 (200 kV), equipped with a Gatan K3 Summit direct detector. Pixel size was 1.219 Å. The movies were captured in super resolution mode with electron dose rate at 10 electrons per pixel per second for 8 s and 0.2 s per frame. Defocus range was −1.0 to −2.5 µm.

Both data sets were processed (Motion correction and CTF estimation) by cryoSPARC (v2.12.4) (*8, 9*). Images were evaluated by inspecting the CTF fitting resolution. Roughly 1000 particles were then manually picked for generating autopicking templates through 2D classification in CryoSPARC. For MV-NLP with 147bp DNA, this yielded 2,648,493 particles across 6,552 micrographs, and these particles were subjected to four iterative rounds of 2D classification in order to discard bad particles. Two ab initio models were then generated, followed by heterogeneous refinement of the nucleosome-like class (34,418 particles). Non-uniform refinement, as well as global and local CTF refinements, were performed to improve the density to the final resolution, as estimated by GFSC. For MV-NLP with 207bp DNA, template-based particle picking yielded ~ 2,131,563 particles across 3,495 micrographs, and these particles were subjected to four iterative rounds of 2D classifications to get best nucleosome particles. Two *ab initio* models were then generated, followed by heterogeneous refinement of the nucleosome like class. Non-uniform refinement with 377,702 particles, were performed to improve the density to the final resolution, as estimated by GFSC.

Initial models were built by fitting MV histone homology models (see below) into the final 3D electron maps in UCSF Chimera, yielding an initial correlation coefficient of 0.7086 (*10*). These models were then iteratively modified and locally refined in COOT (*11*).Molecular dynamics flexible-fitting (MDFF, details below) was then used utilized to simultaneously balance correlation with the EM density and atomic-level energy evaluation (electrostatic configurations and removal of steric overlaps).

**Homology modeling**

Initial homology models of Melbournevirus ORF 369 (H2B-H2A) and ORF 368 (H4-H3) doublets were constructed using SWISSMODEL, and the histone structures from the *Xenopus laevis* nucleosome (PDB 1AOI) were used as a reference ((*12-14*). Using Modeller (v9.20), 10,000 loops of each doublet (H2B-H2A Y152 to S192 and H4-H3 F122 to T203, with sequence numbers identified as in Fig. S1A) were generated (*15*). To eliminate improbable loops within each doublet, clash identification was performed using CPPTRAJ of the Amber MD package (v18), where a cutoff distance of 0.8 Å between potentially over-lapping atoms was used (*16*). Any loops identified by CPPTRAJ without a clash were manually parsed through to eliminate loops that were not sterically overlapping but also not physically relevant, such as those forming "knots" in the histone folds. While several configurations with reasonable qualitative agreement to the EM density were identified (Fig. S5), the loop with the overall best correlation with the density was selected as the initial conformation for further refinement via COOT (*17*), PHENIX (*18*), and MDFF (see below) (*19*). Figures were rendered using Chimera and VMD (*20*).

**Molecular Dynamics Flexible Fitting (MDFF) Protocol**

MDFF simulations were initialized from the PHENIX-refined structure based upon our initial homology modeling. This method improves fitting to EM data by introducing external forces, which bias the structure from its initial conformation into the empirical density. Simulations were conducted using the GPU-enhanced NAMD engine (v2.13) with the CHARMM36 forcefield in an implicit solvent environment (*21, 22*), biasing forces were applied to the protein backbone and heavy atoms of the DNA, in accordance with the level of resolution provided by the density. The simulation timestep was 2 fs, and a total simulation time of 5 ns was sufficient for convergence into the empirical density. Distance- and angle-based restraints of 20 kcal/mol/Å$^2$ and 20 kcal/mol/degree$^2$ were applied to structured regions to prevent over-fitting of the density by breaking of domains with known secondary structure, and dihedral restraints of 50 kcal/mol/degree$^2$ were applied to maintain proper chirality during the biasing process. Over the course of the simulation, correlation between the model and experimental density only showed a modest improvement from 0.761 to 0.766, but significant benefits to residue conformations were observed, such as the removal of drastic $\Phi$ and $\Psi$ orientations or the repositioning of sidechains that were originally used to explain main chain densities due to overfitting by real-space refinements in PHENIX.

**Structural Characterization of the MV-NLP$_{147}$ Model**

Comparisons between MV-NLP$_{147}$ and eNuc$_{147}$ (PDB 3LZ0; (*23*)) were conducted using VMD, and the protein backbone (N, C, and Cα) and DNA sugar ring (C1', C2', C3', C4', and C5') atoms were used for RMSD and center-of-geometry calculations (Table S3). To minimize discrepancies between each system, the eNuc$_{147}$ molecule was least-squares fit to the MV-NLP$_{147}$ using the core α-helices and central 121 bp of DNA as a reference. In this way, RMSD values represent the changes in the global positioning of each constituent, rather than measuring the conformational rearrangement within each individual piece, which was altogether quite low (typically ~1-2 Å). MV-H2B-H2A dimer reorientation, relative to the eukaryotic H2A-H2B dimer, was determined by placing fictitious particles at the geometric centers of the first and last turns of the H2B $\alpha_2$ helix and calculating the dihedral angle describing this imaginary atom arrangement (Fig. S5A). In the eNuc$_{147}$ system, the fictitious atoms describing the C-terminal end of each helix were not located at the final turn of the helix, but rather at the turn within the helix that would yield an analogous length of the H2B $\alpha_2$ helix in MV-NLP$_{147}$, to provide the best direct comparison of the two systems. Larger magnitude values in this measurement correspond to dimers that are angled further away from the dyad-axis, as well as from one another.

Figure S1A

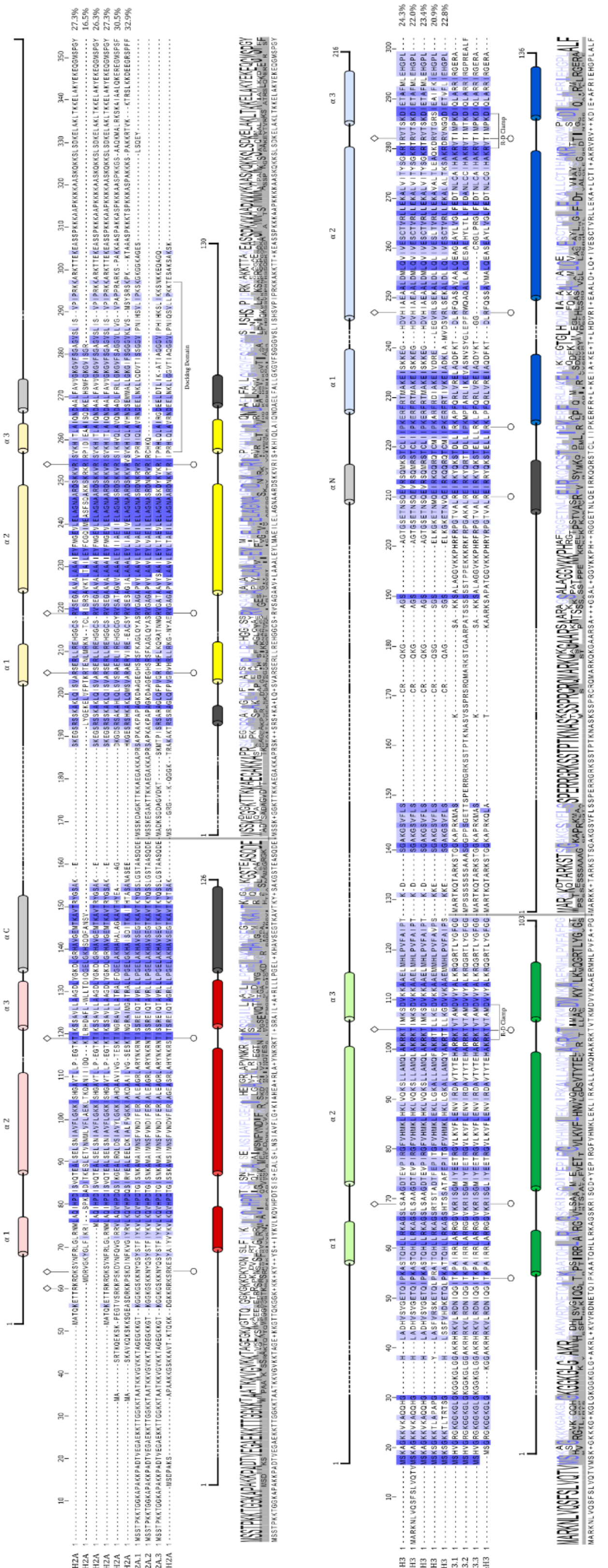

Figure S1 B

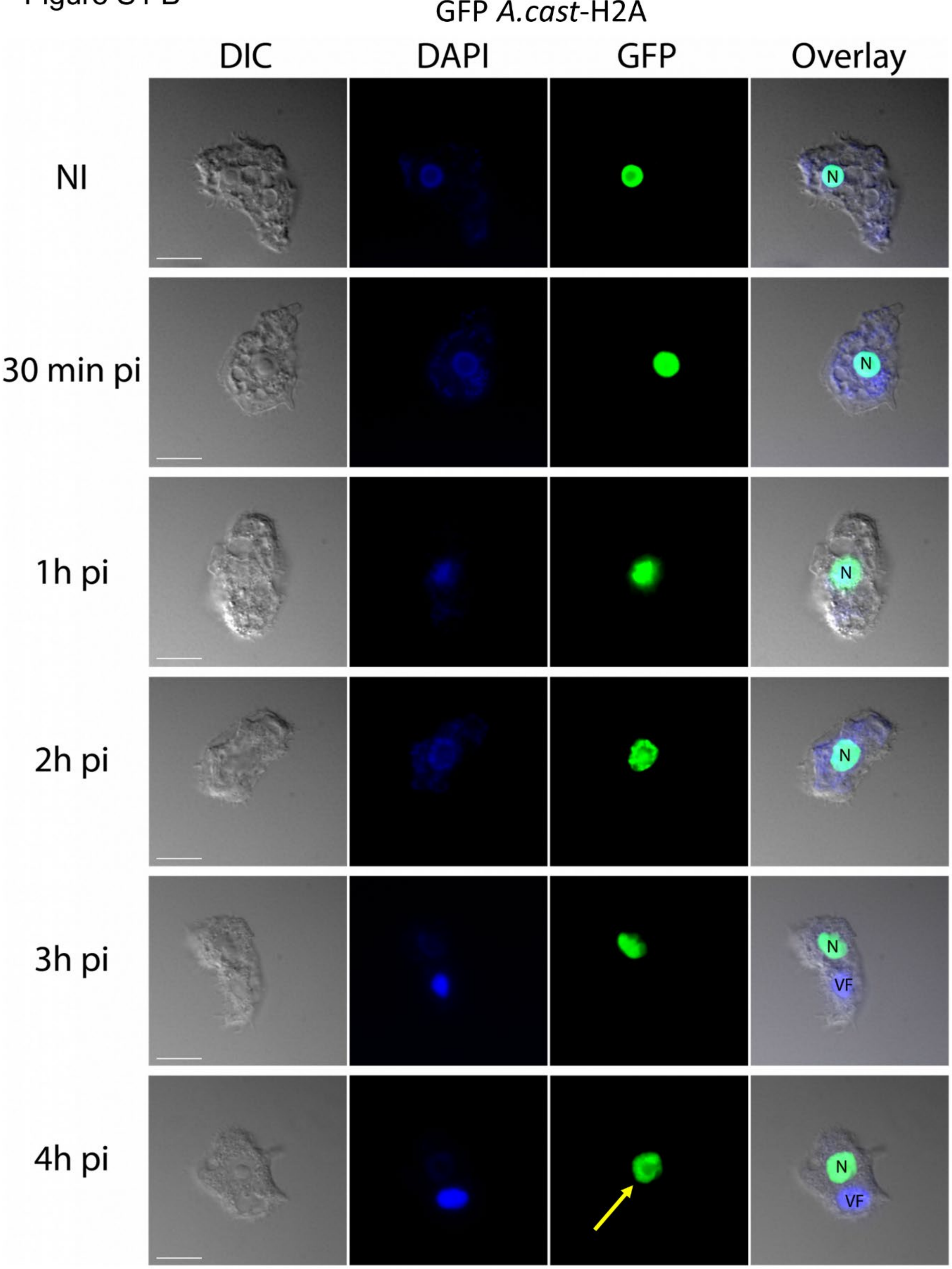

GFP *A.cast*-H2A
DIC
DAPI
GFP
Overlay
NI
30 min pi
1h pi
2h pi
3h pi
4h pi
N
VF

Figure S1 C

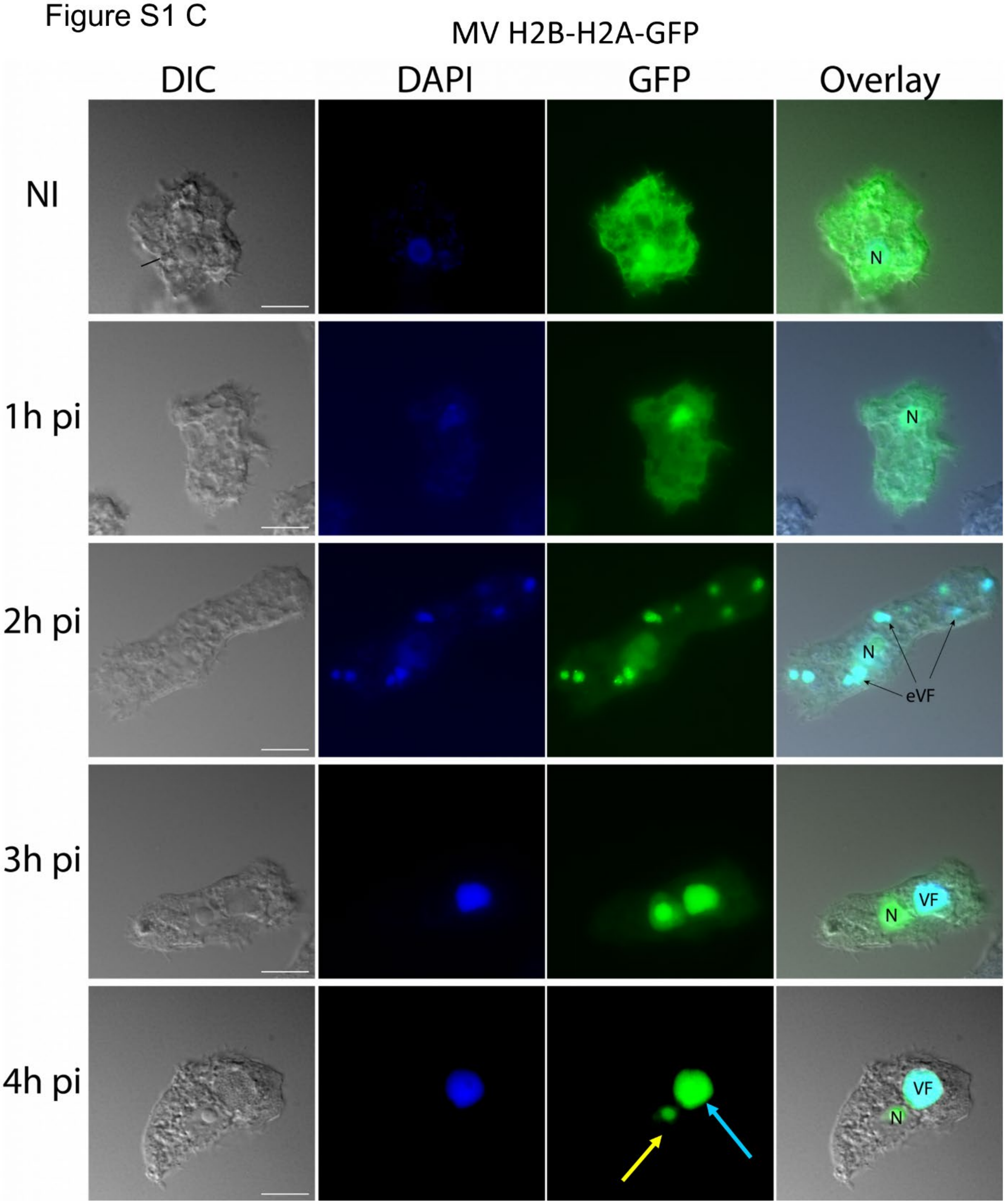
MV H2B-H2A-GFP
DIC
DAPI
GFP
Overlay
NI
N
1h pi
N
2h pi
N
eVF
3h pi
VF
N
4h pi
VF
N

Figure S1 D

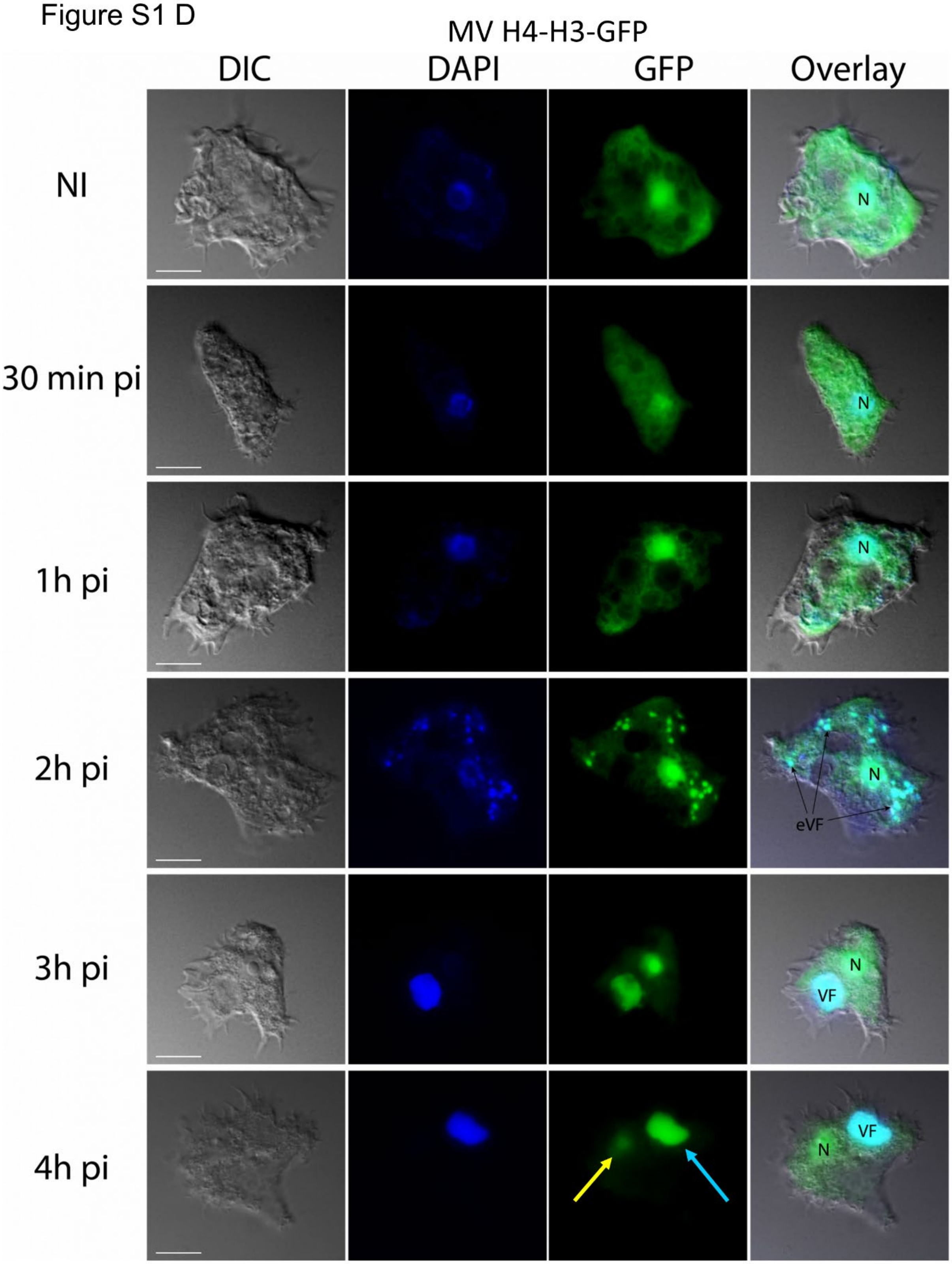
MV H4-H3-GFP
DIC
DAPI
GFP
Overlay
NI
N
30 min pi
N
1h pi
N
2h pi
N
eVF
3h pi
N
VF
4h pi
VF
N

Figure S1 E

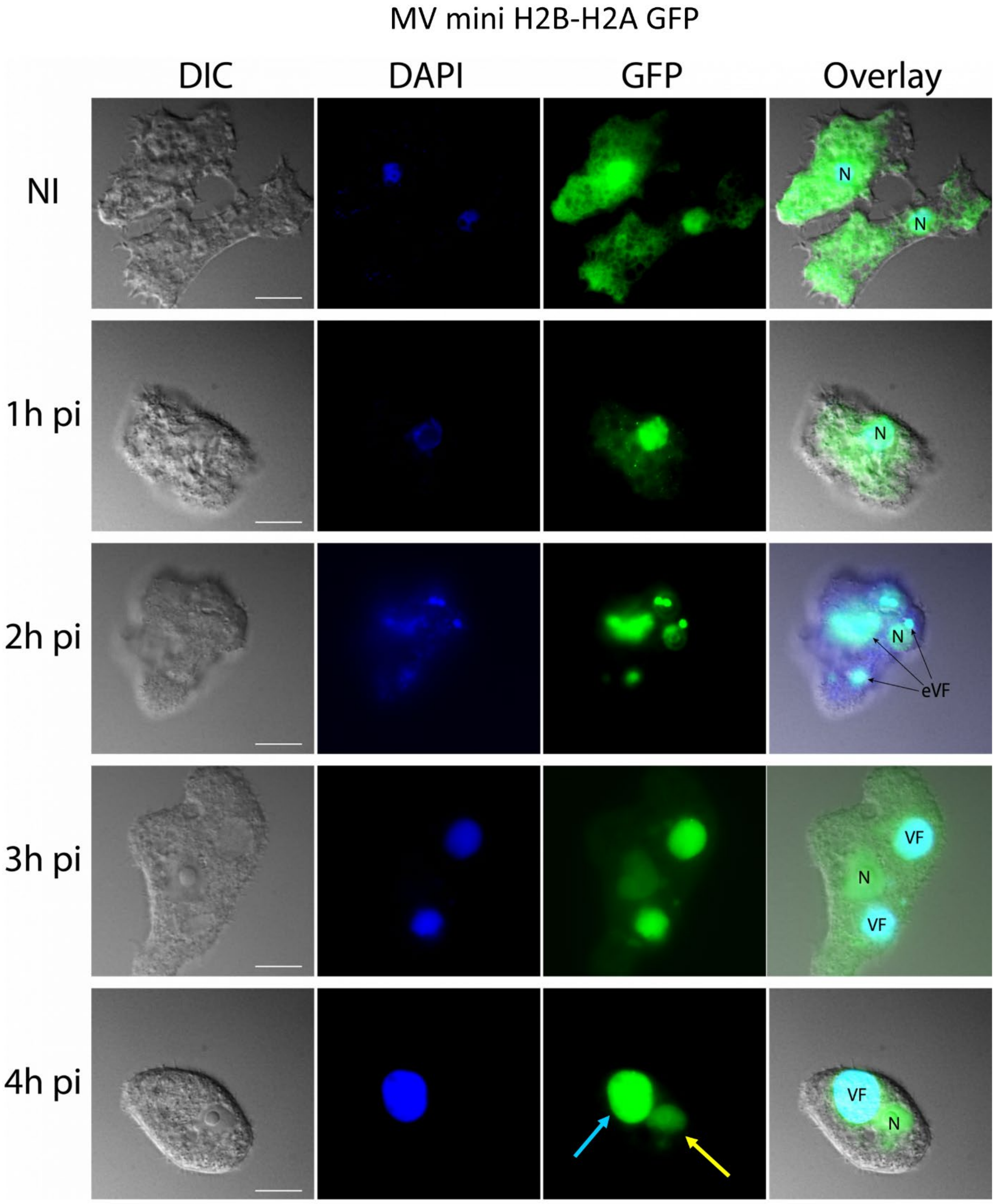
MV mini H2B-H2A GFP
DIC
DAPI
GFP
Overlay
NI
N
N
1h pi
N
2h pi
N
eVF
3h pi
VF
N
VF
4h pi
VF
N

**Fig. S1. Localization of fluorescently labeled Melbournevirus histones in virus-infected Amoeba. (A)** Complete sequence alignment of H2B-H2A (top) and H4-H3 (bottom) from *Marseilleviridae*, the host *Acanthamoeba castellanii*, and *Xenopus laevis.* Predicted α helices of H2B-H2A (light red and yellow) and H4-H3 (light green and blue) *Melbournevirus* histone doublets were generated using HHPRED's Quick 2D prediction web server. Histone dimer pairs H2B-H2A and H4-H3 of *Acanthamoeba castellanii* and *Xenopus laevis* were each aligned against their respective *Marseilleviridae* histone doublets using HHPRED's multiple sequence alignment tool, ClustalΩ. Conservation of each specific residue in each alignment is denoted by blue shading, with greater conservation being represented by darker blue. Known R-D clamp, R-T pairs, and DNA binding residues are indicated for *Xenopus laevis* histone pairs with their conservation within Melbournevirus histones. *Marseilleviridae* histones are 16-33% conserved to *Xenopus laevis* histones (shown on far right of alignment). Known α helices from the histone fold domain in *Xenopus laevis* are shown as dark colored tubes; H2B are red, H2A are yellow, H4 are green, H3 in blue, and additional helices in grey. Logo plot demonstrating residue conservation among the alignments provided by ClustalΩ tool is shown below. Light microscopy fluorescence images (scale bar 10 µm) of *A. castellanii* cells transfected with **B)** GFP-A. castellanii-H2A, **C)** MV-H2B-H2A-GFP, **D)** MV-H4-H3-GFP and **E)** MV-miniH2B-H2A-GFP, non-infected (NI) and infected with Melbournevirus at 30 min, 1h, 2h, 3h and 4h pi. A) GFP-*A. castellanii*-H2A concentrates only in the nucleus (N) of the non-infected cells and all along the infectious cycle. **C)** MV-H2B-H2A-GFP, **D)** MV-H4-H3-GFP and **E)** MV-miniH2B-H2A-GFP are scattered in the entire cell (including the nucleus) in the non-infected cells. Between 1h and 2h pi, the viral histones start accumulating in the early viral factories (eVF). At 4h pi, the fluorescence is predominantly concentrated in the mature viral factory (VF). DAPI staining remains at the nucleus all along the infection but the intense fluorescence in the late VF hides the staining of the nucleus at 4h pi.

### Figure S2

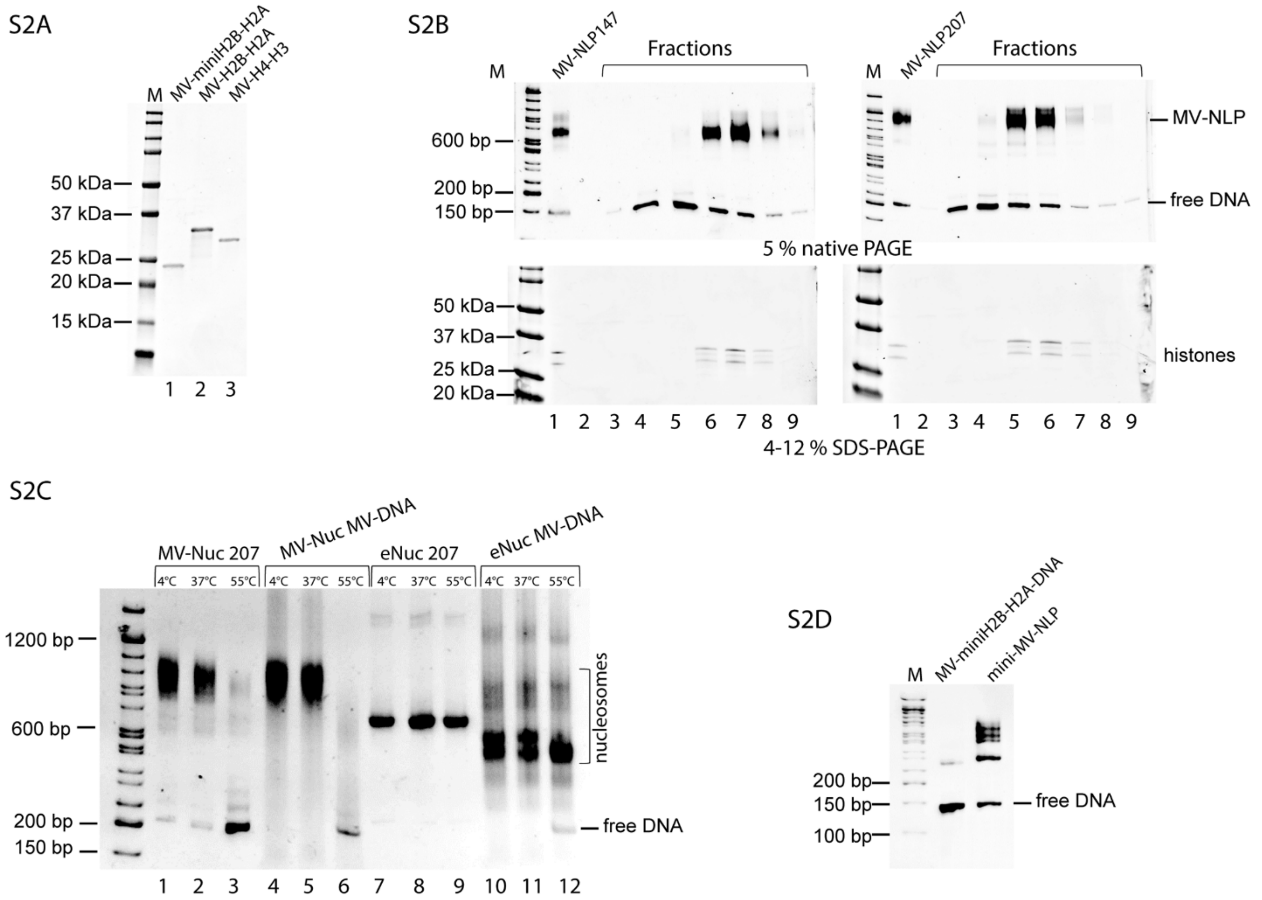

S2A
M
MV-miniH2B-H2A
MV-H2B-H2A
MV-H4-H3
50 kDa
37 kDa
25 kDa
20 kDa
15 kDa
1 2 3
S2B
M
MV-NLP147
Fractions
MV-NLP207
Fractions
MV-NLP
600 bp
200 bp
150 bp
free DNA
5 % native PAGE
50 kDa
37 kDa
25 kDa
20 kDa
histones
1 2 3 4 5 6 7 8 9
1 2 3 4 5 6 7 8 9
4-12 % SDS-PAGE
S2C
MV-Nuc 207
MV-Nuc MV-DNA
eNuc 207
eNuc MV-DNA
4°C 37°C 55°C
1200 bp
600 bp
200 bp
150 bp
nucleosomes
free DNA
1 2 3 4 5 6 7 8 9 10 11 12
S2D
M
MV-miniH2B-H2A-DNA
mini-MV-NLP
200 bp
150 bp
100 bp
free DNA

Figure S2 E, F

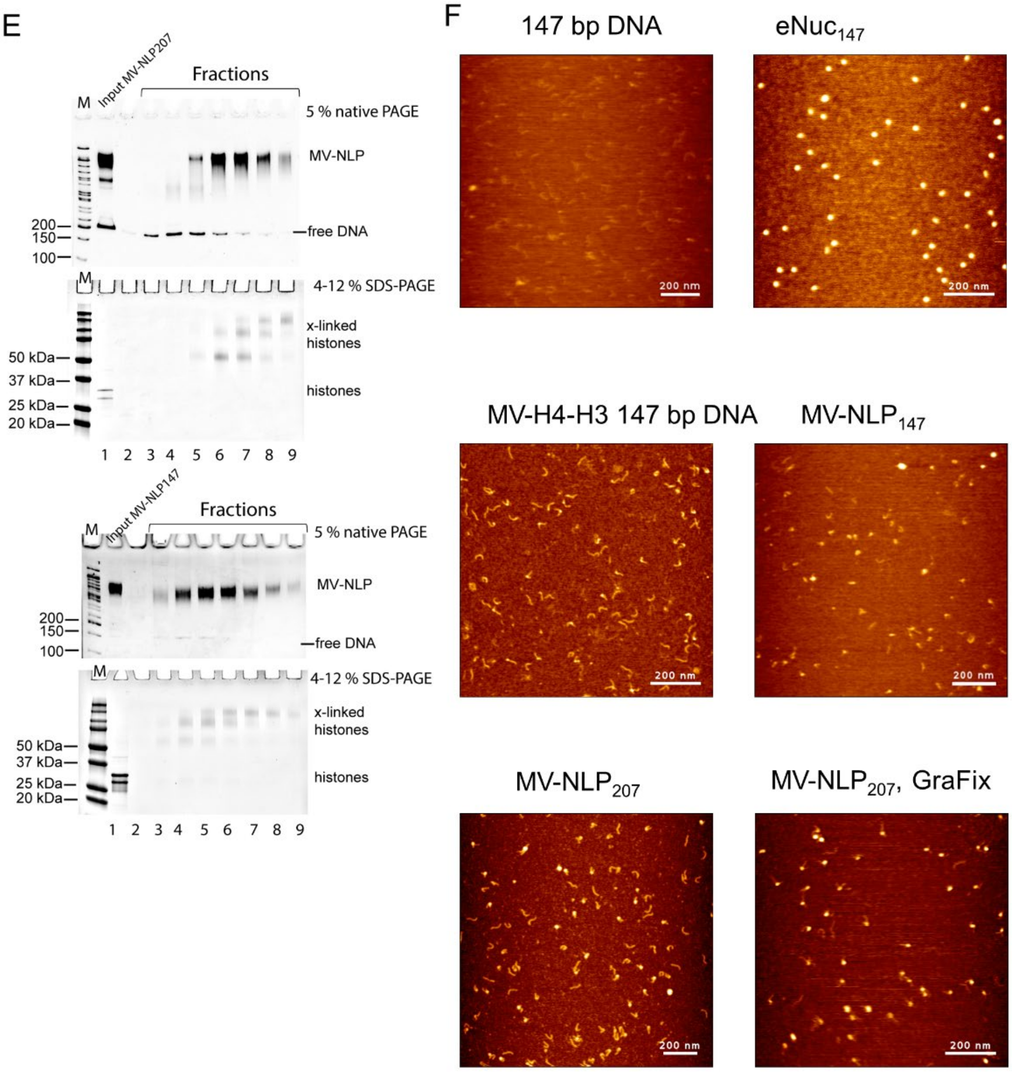

**Fig. S2. MV-histones form nucleosome-like particles (A)** SDS-PAGE of purified Melbournevirus (MV) histone doublets. **(B)** Sucrose gradient sedimentation of MV NLPs with 147 or 207 bp DNA. The compositions of MV NLPs were analyzed by native- and SDS-PAGE. **(C)** MV-NLP and eNuc were reconstituted on Widom '601' DNA and Melbournevirus native DNA, respectively. The MV-NLP and eNuc were heat treated at 37 and 55 ˚C. **(D)** Native PAGE of reconstituted MV mini-NLP (mini H2B-H2A instead of H2B-H2A) with 147 bp DNA. **(E)** GraFix of MV-NLPs with 207 bp DNA (top two panels) and 147 bp DNA (bottom two panels. Native- and SDS- PAGE of the crosslinked MV-NLP fractions representing successful crosslinking, compared to the native MV-NLP input. **(F)** Representative AFM images: Samples were diluted in TCS buffer and applied to APTES coated mica, rinsed with water and imaged on a NanoWizard Sa with a TAP300-GD cantilever. Samples include 147 bp DNA only, $eNuc_{147}$, MV-H4-H3 with 147 bp DNA, $MV\text{-}NLP_{147}$, $MV\text{-}NLP_{207}$ and $MV\text{-}NLP_{207}$ GraFix.

Figure S3

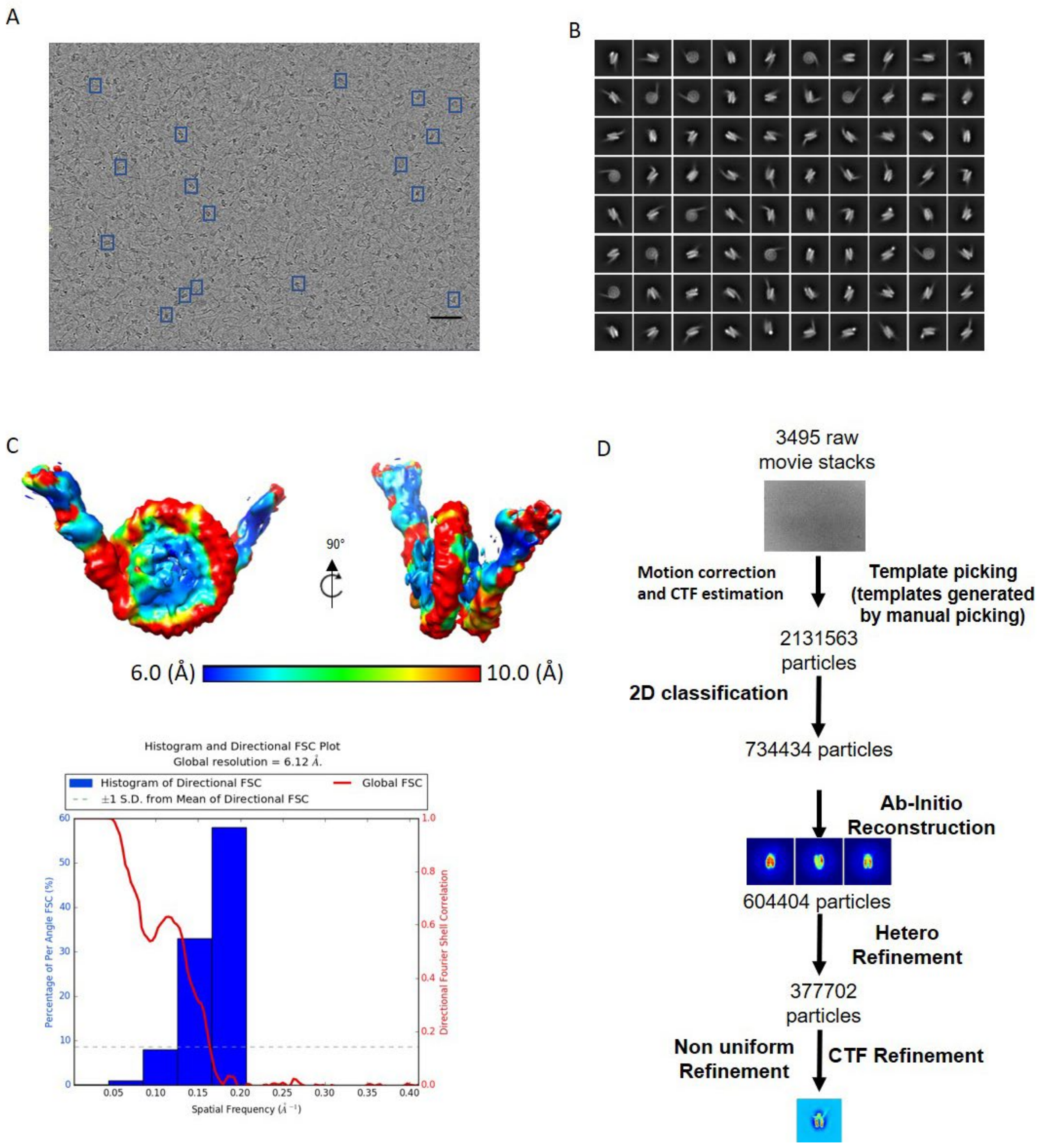

**Fig. S3.** Cryo EM data analysis of MV-NLP$_{207}$. (A) Raw micrograph of the Grafix-ed MV-NLP$_{207}$. Scale bar is 50 nm. (B) 2D class averages generated from the dataset. (C) 3D structure of the MV NLP. Local resolution map and FSC curve. (D) Data processing strategy flow chart.

Figure S4

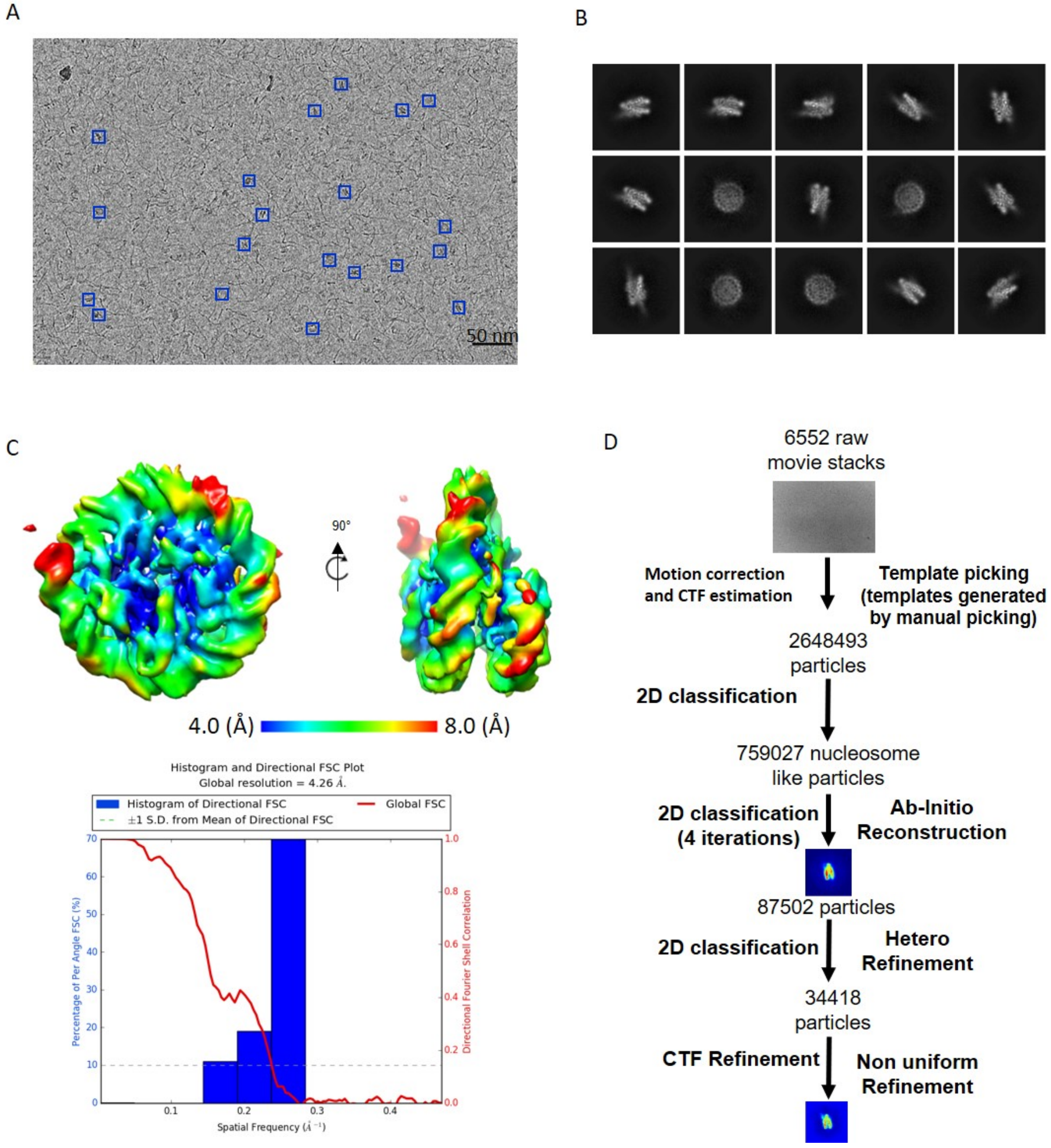

**Fig. S4.** Cryo EM data analysis of MV-NLP$_{147}$. **(A)** Raw micrograph of the GraFix treated MV-NLP$_{147}$. Scale bar is 50 nm. **(B)** 2D class averages generated from the dataset. **(C)** 3D structure of the MV NLP, and local resolution map and FSC curve. **(D)** Data processing strategy flow chart.

Figure S5

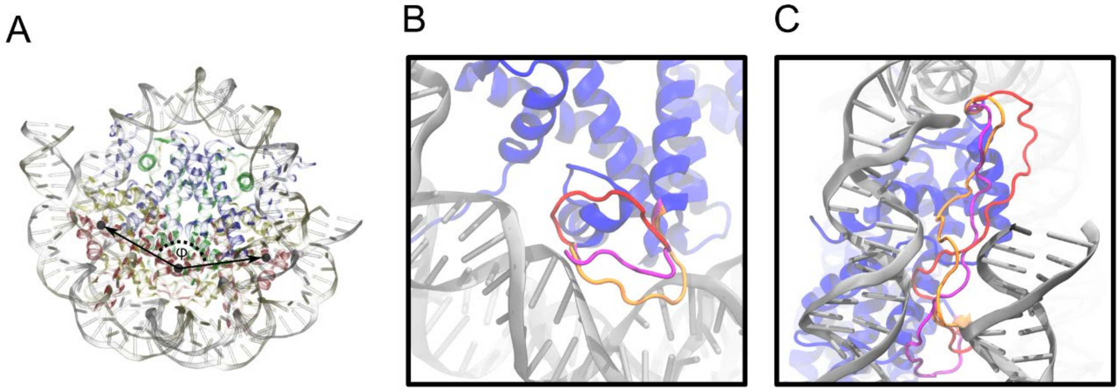

**Fig. S5.** (A) Description of ϕ, the pseudo-dihedral angle measuring the relative orientations of the H2B-H2A subunits, between MV-NLP and eNuc. Larger magnitude values correspond to a dimer arrangement that is angled further away from the dyad axis. Fictitious particle locations are shown as black spheres, and histones (H3 in blue, H4 in green, H2A in yellow, and H2B in red) and DNA (silver and gold ribbons) are semi-transparent to allow for better visibility of each particle's location. **(B)** Overlay of *de novo* loop models with feasible conformations connecting H2B to H2A and **(C)** H4 to H3 in the MV-histone doublets. Loop configurations were generated using Modeller (v9.20) and then placed in the nucleosome context. After manually removing physically non-relevant (knots) conformations and clashes identified with a cutoff distance of 0.8 Å using CPPTRAJ of the Amber MD package (v18), the three best loops are shown in different colors (cross correlation range against the simulated map is 0.614-0.623).

**Table S1. Analysis of AFM data.** Samples were imaged by AFM with heights (nm) of nucleosome like particles (NLP) recorded separately from free DNA. Average heights and standard deviations were calculated from the number of incidences (n). In addition to the average height of the NLP, the amount of free DNA (as a percent of particles on the slide) is a good indication of the stability of MV-NLP. When not prepared by GraFix, NLP heights generally correspond with heights of the MV-Tetrasome (p-values from unpaired t-test), whereas GraFix preparation yields heights in agreement with the $eNuc_{147}$ system.

| | *MV-H4-H3-DNA* ***(MV-tetrasome)*** | | ***eNuc<sub></sub>*** | | | | | | | |
|---|---|---|---|---|---|---|---|---|---|---|
| ***Particle Class*** | **Free DNA** | **MV-Tetrasome** | **Free DNA** | **eNuc** | **Free DNA** | **MV-NLP** | **Free DNA** | **MV-NLP** | **DNA** | **MV-NLP** |
| ***Average Height (nm)*** | 0.60 | 1.35 | 0.61 | 2.09 | 0.50 | 1.21 | 0.78 | 1.55 | 0.64 | 2.25 |
| ***Heights σ (nm)*** | 0.07 | 0.33 | 0.21 | 0.52 | 0.12 | 0.91 | 0.08 | 0.40 | 0.19 | 0.63 |
| ***n*** | 25 | 26 | 2 | 41 | 29 | 46 | 41 | 37 | 31 | 120 |
| ***% Particl.*** | 43.9 | 45.6 | 4.3 | 87.2 | 32.6 | 51.7 | 50.6 | 45.7 | 20.0 | 77.4 |
| ***p-value (MV-Tet)*** | --- | --- | --- | < 0.0001 | --- | 0.4528 | --- | 0.0403 | --- | < 0.0001 |
| ***p-value ($eNuc_{147}$)*** | --- | < 0.0001 | --- | --- | --- | < 0.0001 | --- | < 0.0001 | --- | 0.1452 |

**Table S2. Summary of cryoEM data collection and refinement**

| | $MV\text{-}NLP_{147}$ (EMDB-xxxx) (PDB xxxx) | $MV\text{-}NLP_{207}$ |
|---|---|---|
| **Data collection and processing** | | |
| Magnification | 64,000 | 29000 |
| Voltage (kV) | 300 | 200 |
| Electron exposure (e–/Å$^2$) | 50 | 80 |
| Defocus range (µm) | 0.8-2.0 | 1.0-2.5 |
| Pixel size (Å) | 1.065 | 1.219 |
| Symmetry imposed | C1 | C1 |
| Initial particle images (no.) | 2,648,493 | 2,131,563 |
| Final particle images (no.) | 34,418 | 377,702 |
| Map resolution (Å) | 4.17 | 6.1 |
| FSC threshold | 0.143 | 0.143 |
| Map resolution range (Å) | 3.9-6.9 | 6.0-10.0 |
| **Refinement** | | |
| Initial model used (PDB code) | 1AOI, 3LZ0 | |
| Model resolution (Å) | 4.1 | |
| FSC threshold | 0.143 | |
| Model resolution range (Å) | 4.1-7.3 | |
| Map sharpening *B* factor (Å$^2$) | | |
| Model composition | | |
| Non-hydrogen atoms | 11125 | |
| Protein residues | 789 | |
| Nucleotide | 250 | |
| Ligands | 0 | |
| *B* factors (Å$^2$) | | |
| Protein | n/a | |
| Nucleotide | n/a | |
| Ligand | n/a | |
| R.m.s. deviations | | |
| Bond lengths (Å) | 0.015 | |
| Bond angles (°) | 1.805 | |
| Validation | | |
| MolProbity score | 1.50 | |
| Clashscore | 0.00 | |
| Poor rotamers (%) | 1.5 | |
| Ramachandran plot | | |
| Favored (%) | 93.67 | |
| Allowed (%) | 5.06 | |
| Disallowed (%) | 1.27 | |

**Table S3: Comparison of MV-NLP$_{147}$ with eNuc$_{147}$.** “Global” root-mean-squared deviation (RMSD) values represent changes in positioning throughout the nucleosome-like complex and are calculated for each domain after least-squares fitting the cores of the MV-NLP$_{147}$ and eNuc$_{147}$ complexes, whereas “local” RMSDs are values calculated when least-squares fitting individual domains prior to calculation and thereby represent the degree of internal re-ordering of the domain. Distances are calculated as the separation of geometric centers of backbone positions (C, $C_\alpha$, and N atoms), except for the H2A L1 Loops “closest distance”, which defines the smallest distance between heavy atoms in the two moieties. The H2B $\alpha_C$ orientation angle is calculated around the pseudo-dihedral defined in Figure S5A.

| **RMSD Values (Å)** | **Global** | **Local** |
|---|---|---|
| Particle Core | 2.7 | --- |
| DNA (central 120 bp) | 3.1 | 3.1 |
| Octamer Fold | 2.3 | 2.3 |
| H4-H3 Tetramer | 2.1 | 1.2 |
| H3 $\alpha_N$-helix (Chain A) | 2.3 | 0.3 |
| H3 $\alpha_N$-helix (Chain E) | 2.2 | 0.3 |
| H2B-H2A Dimers (both) | 2.5 | 2.1 |
| H2B-H2A Dimer (Chain C,D) | 2.5 | 1.6 |
| H2B-H2A Dimer (Chain G,H) | 2.5 | 1.6 |
| H2B $\alpha_C$-helix (chain D) | 2.6 | 0.5 |
| H2B $\alpha_C$-helix (chain H) | 3.4 | 0.9 |
| H2A Docking Domain (Chain C) | 8.5 | 3.9 |
| H2A Docking Domain (Chain G) | 6.2 | 3.3 |
| **Separation Distances (Å)** | **eNuc$_{147}$** | **MV-NLP$_{147}$** |
| H4-H3 Dimers | 32.6 | 33.3 |
| H3-H3’ 4HB Interface | 10.1 | 10.4 |
| H2B-H2A Dimers | 36.3 | 38.6 |
| H2B $\alpha_C$-helices | 49.1 | 52.2 |
| H2A L1 Loops (centers) | 7.2 | 9.7 |
| H2A L1 Loops (closest distance) | 2.9 | 5.3 |
| **Orientation Angle (°)** | **eNuc$_{147}$** | **MV-NLP$_{147}$** |
| H2B $\alpha_2$-helices | 135.4 | 143.9 |

**Movie S1.**

CryoEM electron density map of MV-NLP$_{207}$ reveals its overall nucleosome-like shape and defined extended linker DNA, which includes a vertical and horizontal dyad axis rotation of MV-NLP$_{207}$ with a density flare.

**Movie S2**

CryoEM electron density map of MV-NLP$_{147}$ reveals its overall nucleosome-like shape, which includes a vertical and horizontal dyad axis rotation of MV-NLP$_{147}$ with a density flare.

**Movie S3**

Overview of MV-NLP$_{147}$ structure (model to map, Fig 3A), including a highlight of doublet connectors and H4 N-term domain (Fig. 4). The equivalent regions of H3, H4, H2A, and H2B are shown in blue, green, yellow, and red, respectively.

1. L. Zimmermann *et al.*, A Completely Reimplemented MPI Bioinformatics Toolkit with a New HHpred Server at its Core. *J Mol Biol* **430**, 2237-2243 (2018).
2. E. Bateman, Expression plasmids and production of EGFP in stably transfected Acanthamoeba. *Protein Expr Purif* **70**, 95-100 (2010).
3. P. N. Dyer *et al.*, Reconstitution of nucleosome core particles from recombinant histones and DNA. *Methods Enzymol* **375**, 23-44 (2004).
4. B. Demeler *et al.*, Characterization of size, anisotropy, and density heterogeneity of nanoparticles by sedimentation velocity. *Anal Chem* **86**, 7688-7695 (2014).
5. G. Gorbet *et al.*, A parametrically constrained optimization method for fitting sedimentation velocity experiments. *Biophys J* **106**, 1741-1750 (2014).
6. E. Brookes, B. Demeler, M. Rocco, Developments in the US-SOMO bead modeling suite: new features in the direct residue-to-bead method, improved grid routines, and influence of accessible surface area screening. *Macromol Biosci* **10**, 746-753 (2010).
7. G. B. Edwards, U. M. Muthurajan, S. Bowerman, K. Luger, Analytical Ultracentrifugation (AUC): An Overview of the Application of Fluorescence and Absorbance AUC to the Study of Biological Macromolecules. *Curr Protoc Mol Biol* **133**, e131 (2020).
8. S. Q. Zheng *et al.*, MotionCor2: anisotropic correction of beam-induced motion for improved cryo-electron microscopy. *Nat Methods* **14**, 331-332 (2017).
9. K. Zhang, Gctf: Real-time CTF determination and correction. *J Struct Biol* **193**, 1-12 (2016).
10. E. F. Pettersen *et al.*, UCSF Chimera--a visualization system for exploratory research and analysis. *J Comput Chem* **25**, 1605-1612 (2004).
11. P. Emsley, K. Cowtan, Coot: model-building tools for molecular graphics. *Acta Crystallogr D Biol Crystallogr* **60**, 2126-2132 (2004).
12. A. Waterhouse *et al.*, SWISS-MODEL: homology modelling of protein structures and complexes. *Nucleic Acids Res* **46**, W296-W303 (2018).
13. S. Bienert *et al.*, The SWISS-MODEL Repository-new features and functionality. *Nucleic Acids Res* **45**, D313-D319 (2017).

14. N. Guex, M. C. Peitsch, T. Schwede, Automated comparative protein structure modeling with SWISS-MODEL and Swiss-PdbViewer: a historical perspective. *Electrophoresis* **30 Suppl 1**, S162-173 (2009).
15. A. Sali, T. L. Blundell, Comparative protein modelling by satisfaction of spatial restraints. *J Mol Biol* **234**, 779-815 (1993).
16. D. R. Roe, T. E. Cheatham, 3rd, PTRAJ and CPPTRAJ: Software for Processing and Analysis of Molecular Dynamics Trajectory Data. *J Chem Theory Comput* **9**, 3084-3095 (2013).
17. P. Emsley, B. Lohkamp, W. G. Scott, K. Cowtan, Features and development of Coot. *Acta Crystallogr D Biol Crystallogr* **66**, 486-501 (2010).
18. P. D. Adams *et al.*, PHENIX: a comprehensive Python-based system for macromolecular structure solution. *Acta Crystallogr D Biol Crystallogr* **66**, 213-221 (2010).
19. R. McGreevy, I. Teo, A. Singharoy, K. Schulten, Advances in the molecular dynamics flexible fitting method for cryo-EM modeling. *Methods* **100**, 50-60 (2016).
20. W. Humphrey, A. Dalke, K. Schulten, VMD: visual molecular dynamics. *J Mol Graph* **14**, 33-38, 27-38 (1996).
21. K. Y. Chan, L. G. Trabuco, E. Schreiner, K. Schulten, Cryo-electron microscopy modeling by the molecular dynamics flexible fitting method. *Biopolymers* **97**, 678-686 (2012).
22. J. Huang, A. D. MacKerell, Jr., CHARMM36 all-atom additive protein force field: validation based on comparison to NMR data. *J Comput Chem* **34**, 2135-2145 (2013).
23. D. Vasudevan, E. Y. D. Chua, C. A. Davey, Crystal structures of nucleosome core particles containing the '601' strong positioning sequence. *J Mol Biol* **403**, 1-10 (2010).
